# Genome-wide Identification of Type VI Secretion System Effectors via Machine Learning Uncovers Three New Bacterial Toxins

**DOI:** 10.64898/2026.09.28.754971

**Authors:** Avital Akerman-Arad, Aleks Danov, Yuval Chausho, Aya Friedman, Shai Benedict, Maor Shalom, Tal Fisher, Rina Fraenkel, Gideon Mamou, Yaara Oppenheimer-Shaanan, Netanel Tzarum, Asaf Levy

## Abstract

The type VI secretion system (T6SS) is a molecular harpoon used by 19% of sequenced bacteria to inject toxic proteins into microbial competitors or host cells. Despite its widespread distribution, many T6SS effectors (T6Es) remain unidentified in bacterial genomes. Here, we developed an XGBoost classifier integrating 42 genetic and biochemical features to identify novel T6Es. Applying our model to 2,559 T6SS-encoding bacterial genomes revealed 19,958 T6E candidates. We validated our computational predictions and confirmed that three effector candidates, having new 3D structures, demonstrated cytotoxicity to *Escherichia coli* and/or *Saccharomyces cerevisiae*, mostly in the bacterial periplasm. For two of these, cognate immunity proteins neutralizing cytotoxicity have been validated as well. T6EC11, a new peptidoglycan degrading effector, quickly leads to spherical *E. coli*, with inner membrane separation, and gradual peptidoglycan degradation in a unipolar manner, demonstrating a moon-like structural phenotype. T6EC12, a predicted cargo effector that is toxic to bacteria and yeast, carries a conserved zinc-binding motif. Mutations in the predicted catalytic center markedly reduced toxicity, suggesting that T6EC12 represents a novel metalloprotease T6E. Our study expands the known T6E repertoire and provides thousands of predictions of potential effectors in most T6SS-encoding bacteria.

## Introduction

Bacteria often live in dense mixed polymicrobial communities where they compete fiercely for space and resources. To gain a competitive advantage, many species have evolved secretion systems capable of killing neighboring cells or infecting host organisms. Among these, numerous Gram-negative bacteria have repurposed phage tail-like structures into potent molecular weapons, giving rise to the Type VI Secretion System (T6SS). T6SS function as contractile, harpoon-like nanomachines that deliver toxic effector proteins (T6Es) directly into neighboring cells in a contact-dependent manner, killing or inhibiting competitors (Pukatzki *et al*, 2007). Found in 19% of sequenced bacteria, mostly host-associated pathogens (Geller *et al*, 2021; Unni *et al*, 2022), T6SSs deliver T6Es to diverse cellular targets, mostly Gram-negative bacterial competitors, but also into eukaryotic host cells and fungi (MacIntyre *et al*, 2010; Ray *et al*, 2017). The system’s widespread distribution among established pathogens (Rehman *et al*, 2026; Geller *et al*, 2021), combined with experimental evidence demonstrating its crucial role in virulence and colonization across multiple species and habitats, underscores its significance in bacterial pathogenesis (Pukatzki *et al*, 2006; Unni *et al*, 2022).

Structurally, the T6SS comprises 13 conserved components (TssA-M) typically encoded in an operon (Boyer *et al*, 2009). The puncturing apparatus consists of stacked Hcp hexamers forming a tube, capped by a VgrG trimer spike and PAAR protein tip. Upon activation, rapid sheath contraction propels this structure into target cells, delivering T6Es (Kudryashev *et al*, 2015). T6Es reach their targets either as evolved domains fused to Hcp, VgrG, or PAAR domains, or as cargo proteins loaded into the Hcp tube or onto the VgrG spike via chaperones (Wang *et al*, 2023). These T6Es exhibit remarkable enzymatic diversity, targeting membranes (González-Magaña *et al*, 2022; Nicastro *et al*, 2026), cell walls (Whitney *et al*, 2013; Le *et al*, 2021), nucleic acids (Jana *et al*, 2019; Hespanhol *et al*, 2022), or other molecules(Nolan *et al*, 2021) to disrupt essential cellular functions. T6Es can carry one or two toxic domains (Fridman *et al*, 2025) and harbor characteristic motifs and domains, e.g. N-terminal PAAR domains, MIX motifs, and FIX domains (Salomon *et al*, 2014; Jana *et al*, 2019), that aid in their identification (Salomon *et al*, 2014) but importantly T6E proteins lack a characteristic secretion signal. To prevent self-intoxication by antibacterial T6Es, bacteria encode short immunity proteins adjacent to their corresponding T6E genes (Dong *et al*, 2013) although defense independent of immunity proteins has also been reported (Hersch *et al*, 2020a). Recent studies have revealed that T6SS effector repertoires are far more diverse than initially anticipated, with rapid evolutionary turnover driven by interbacterial antagonism (Habich *et al*, 2025).

The identification and characterization of novel T6Es is crucial for understanding microbial interactions (Verster *et al*, 2017) and for biotechnological applications (Mok *et al*, 2020). These toxins represent natural antibacterial compounds that could serve as templates for new antibiotics to combat antimicrobial resistance (Nachmias *et al*, 2024), while also providing insights into bacterial competition (Koskiniemi *et al*, 2013), virulence mechanisms (Jiang *et al*, 2024), and niche adaptation (Shidore *et al*, 2012). Despite their importance, identifying new T6Es is challenging due to their diversity and the absence of characteristic signal peptides (Lewis *et al*, 2019; Liang *et al*, 2015).

These challenges have prompted the development of various detection methods, including experimental screens (Dong *et al*, 2013; Nolan *et al*, 2021), computational approaches such as 3D protein modeling to predict enzymatic activity (Geller *et al*, 2024), comparative genomics (Fridman *et al*, 2020) and sequence homology analysis to detect signature domains (Salomon *et al*, 2014; Jana *et al*, 2019). Machine learning approaches have also been developed to analyze amino acid composition, evolutionary patterns, and physicochemical properties (Sen *et al*, 2019; Jiaweiwang *et al*, 2018; Hu *et al*, 2025). While valuable, these methods have not fully utilized the comprehensive genomic signatures of known T6Es. Moreover, importantly, these methods have not been assessed empirically by experimental validation of new T6Es. The absence of model validation is critical as often there is a dramatic discrepancy between prediction accuracy based on computational cross-validation and experimental validation. This discrepancy may result from over-fitting.

To address this limitation, we developed a machine learning algorithm integrating 42 significant genomic and physicochemical features, enabling the identification of previously undiscovered 19,958 T6Es candidates across all sequenced T6SS-encoding genomes. We observed that T6Es tend to be longer, more hydrophilic, charged, and ordered, and to reside next to or fused to T6SS genes much more than non-T6Es. We then tested eight predicted candidates T6ECs (Type VI effector candidate) for their ability to kill prokaryotic or eukaryotic cells, discovering three novel T6Es that are toxic to *E. coli*, *S. cerevisiae,* or both: T6EC11, T6EC12, and T6EC17 encoded by three diverse environmental bacteria. For all three confirmed effectors, cognate immunity proteins were predicted, two of which were experimentally validated. We followed the phenotypic change occurring upon expression of these effector proteins in *E. coli*. Interestingly, T6EC11, a toxin that was active only in the periplasm, led to cell rounding which is a whole mark of peptidoglycan degradation. However, by following the degradation process we observed inner membrane separation and an unexpected polar accumulation of the peptidoglycan prior to quick cell death. Due to this phenotype, we termed T6EC11 “moon toxin” which resembles the resulting cell shape during intoxication. Purified T6EC11 directly degraded peptidoglycan. In addition, structural characterization of T6EC12 identified a conserved HExxH-containing catalytic center embedded within a previously uncharacterized two-domain scaffold. Mutational analysis demonstrated that residues comprising the predicted catalytic center are required for full toxin activity, suggesting that T6EC12 represents a previously unrecognized metalloprotease T6E. We provide all T6E predictions across the bacterial kingdom for future research by the scientific community.

## Results

### Data collection and feature performances

To create the T6E classifier we first constructed a training dataset. We formed a negative set of non-T6E genes and proteins retrieved from the Integrated Microbial Genomes (IMG) (Markowitz *et al*, 2012). This set included T6SS core genes, random non-toxin genes from T6SS genomes, and effectors from different secretion systems (T1SS-T4SS) (Materials and Methods). The positive dataset was constructed from both experimentally validated T6Es and bioinformatically predicted effectors from the Secret6 database (Li *et al*, 2015). Following the removal of highly similar sequences (>80% sequence identity), the final dataset comprised 634 non-redundant proteins in the positive set and 3,988 in the negative set (Table EV1). A total of 42 features were extracted for each protein in the dataset, encompassing both biochemical properties and the genetic context of their encoding genes, including various genomic annotations, as detailed in the Materials and Methods section.

To evaluate the suitability of the extracted features for T6Es classification, we assessed their ability to distinguish T6Es from non-T6Es. Informative features yielded statistically significant separation between the two groups.

### Genomic and genetic features characterizing T6Es

Analysis of the genomic context and domain presence revealed notable differences between T6Es and non-T6Es, although no single feature provided complete separation (Fig. 1). T6Es genes were located closer to core T6SS machinery than non-T6Es (66% within up to 15 genes away from core components compared to 15% of non-T6SEs; Fig. 1A), illustrating that most T6Es reside within T6SS operons. The inclusion of T6SS core genes within the negative set (to prevent their future classification as T6Es) contributed to the relatively high value (25%) within the non-T6Es. This observation aligns with previous studies documenting "Super-orphan" toxins (Levy *et al*, 2017), which can arise through horizontal gene transfer (Salomon *et al*, 2015). Surprisingly, only 3% of the positive set were located adjacent to genes carrying immunity Pfam (Mistry *et al*, 2021) domains (Fig. S1A), possibly reflecting the existence of novel, uncharacterized immunity genes.

**Figure 1.**
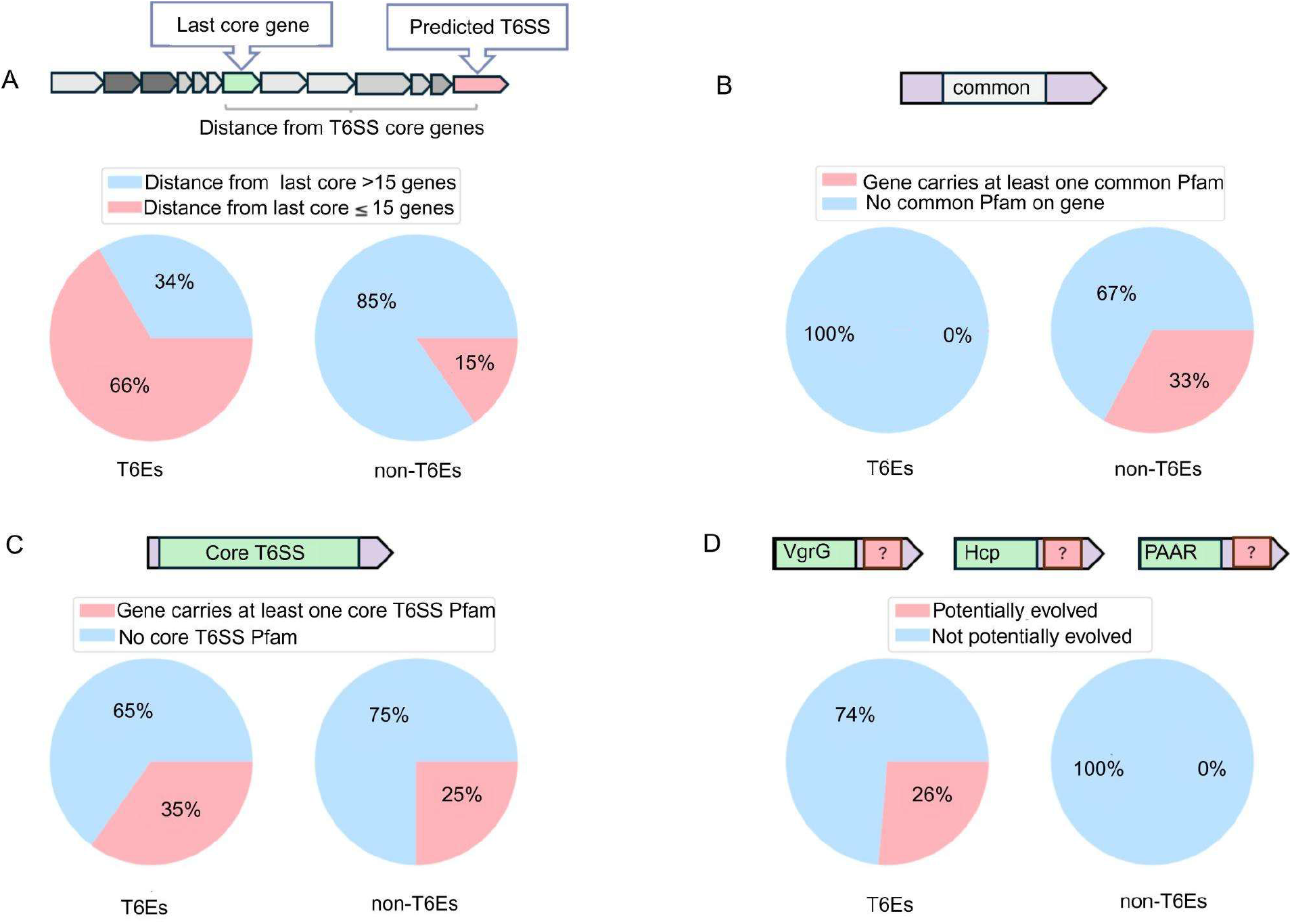
Genomic and genetic features that differentiate between T6Es and the non-T6Es genes. **A.** Gene distance from the closest core T6SS gene. Note that non-T6Es also contain structural T6SS genes leading to their proximity to other core T6SS genes. **B.** Presence of common Pfam domains within the protein. The distribution of most prevalent Pfam domains within our database contain genomes with T6SS (T6SS+), such as ABC transporter and helix-turn-helix domains. These are the top 25% in terms of Pfam prevalence across T6SS+ genes. **C.** T6SS core domain presence. Core T6SS domains, such as VgrG or Hcp, are key components of the T6SS machinery. **D.** Potentially evolved genes. Genes that contain a T6SS structural domain (Hcp, PAAR, or VgrG) in the N-termini regions and have unannotated C-terminal tails of at least 100 amino acids are classified as potentially evolved toxins.

Regarding domain presence, non-T6Es more frequently contained common domains (33% in non-T6Es, vs. 0% in T6Es; Fig. 1B), signal peptides (8% vs. 0%; Fig. S1B), and mobile elements (2% vs. 0%; Fig. S1C). In contrast, T6Es were enriched for T6SS core domains (35% vs. 26%; Fig. 1C) and evolved effector signatures (26% vs. 0%; Fig. 1D), namely VgrG/PaaR/Hcp domains with a C-terminal protein extension. T6Es also carried known toxin domains, eukaryotic-like domains, and glycine zippers more frequently than non-T6Es ( Fig. S1D-F).

Several Clusters of Orthologous Genes (COG) functional categories exhibited significant differences between T6Es and non-T6Es (p<0.05). The most striking enrichment was in three categories in the T6E set: “X” (Mobilome: prophages, transposons), “R” (General function prediction only), and “P” (Inorganic ion transport and metabolism), which showed a significantly higher density for T6Es than non-T6Es (p<0.001). In contrast, the majority of other functional categories had a higher prevalence in non-T6Es (Fig. S2).

Most of the binary features demonstrated limited discriminative power and did not provide a clear separation between the groups. However, many of the examined features showed some degree of separation, suggesting that their integration with each other and other features, such as physicochemical features (Fig. S3), could enhance the performance of a machine learning model in accurately classifying T6Es.

### Physicochemical features characterizing T6Es

To evaluate the potential of various physicochemical features for classifying T6Es, we assessed the discriminatory power of each property using the Kolmogorov-Smirnov (KS) test (Adeodato & Melo, 2016). This analysis revealed several informative features, including mean hydrophobicity, protein length and mean order-promoting amino acids (Materials and Methods).

Mean hydrophobicity demonstrated the strongest discriminatory power (Fig. 2A, KS=0.43, P-value=1.18e-100), with T6Es showing a more hydrophilic tendency, likely due to their constraint to be produced and function in the cytoplasm or periplasm, and often carried inside the T6SS lumen. Protein length distributions also showed marked differences (Fig. 2B, KS=0.49, P-value=1.21e-128). T6Es displayed a multimodal distribution with distinct peaks at 150-400, 700, and 1500 amino acids, while non-T6Es exhibited a unimodal distribution centered between 100-500 amino acids. Long T6Es include the Rhs effectors which carry multiple Rhs repeats (Günther *et al*, 2022). T6Es contain more order promoting residues (I, L, M, V, F, Y, and C residues promote protein folding (Campen *et al*, 2008)) than non-T6Es (Fig. 3C, KS=0.35, P-value=4.33e-67). Mean aromaticity (Fig. 3D, KS=0.27, P-value=1.89e-38) and mean disorder (Fig S3A, KS=0.2, P-value=2.37e-20) demonstrated moderate discriminatory power, while mean molecular weight (Fig S3C, KS=0.12, P-value=1.83e-7) and mean charge (Fig S3B, KS=0.11, P-value=4.59e-07) showed weaker separation between the groups, despite statistical significance.

**Figure 2.**
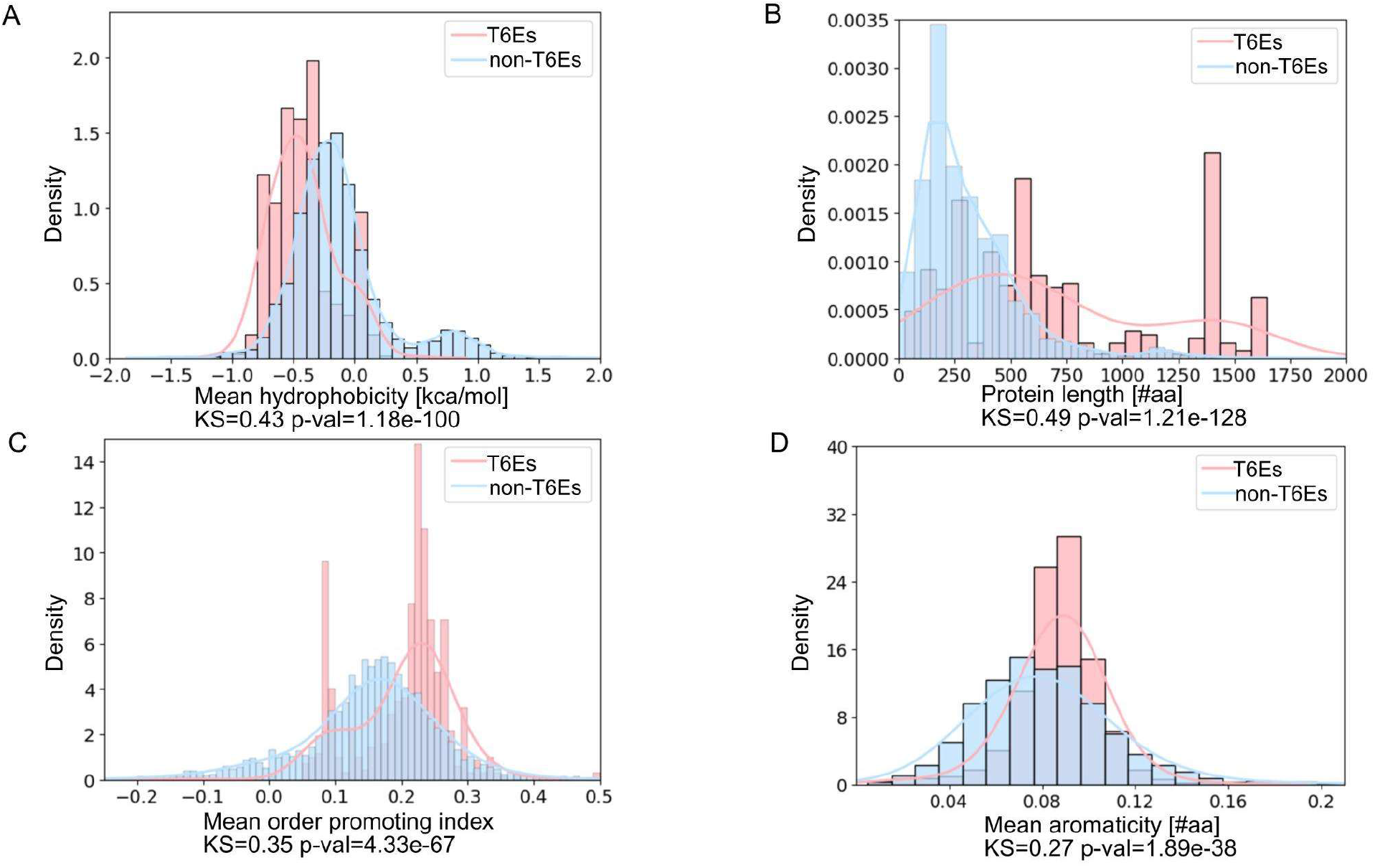
Biochemical features separating between T6Es and non-T6Es. **A.** Protein mean hydrophobicity distribution. **B.** Protein length distribution. **C.** Histogram of protein sequence mean order index. **D.** Mean number of aromatic amino acids within the proteins.

**Figure 3.**
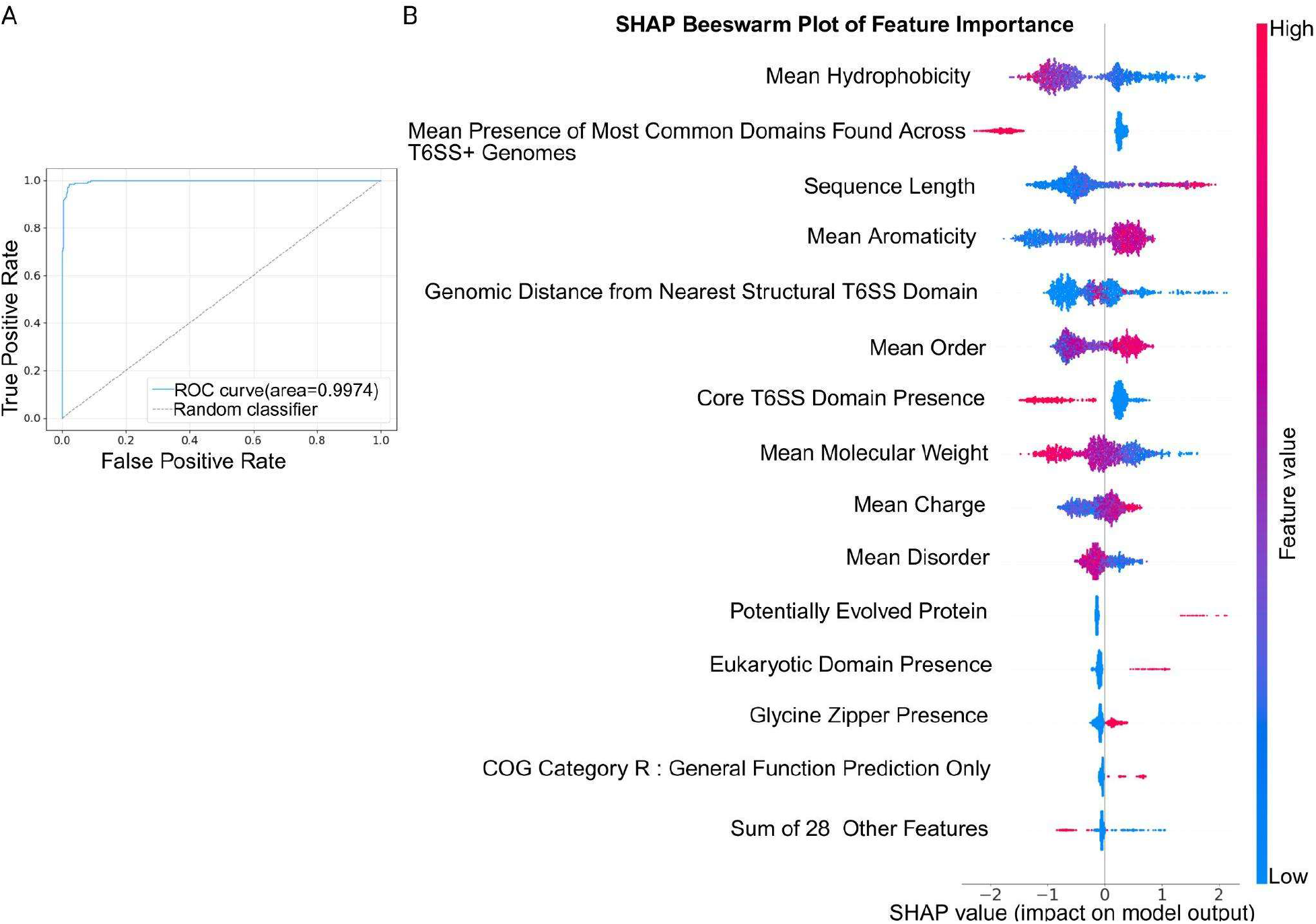
Performance of the T6E XGBoost classification model and its features. **A.** The ROC curve of the classifier. The curve shows the performance of five different models trained on 80% of the data during a 5-fold cross-validation process. **B.** SHapley Additive exPlanations (SHAP) values for each feature of the classification algorithm for T6Es. Each dot represents a single protein from the training set feature matrix. The values are grouped by features and colored by feature values. The red points are high values of the feature, and the blue points are low values. The X-axis represents the SHAP value. Accordingly, the distance from the last core gene and the mean hydrophobicity have the highest contribution to the model.The full feature importance is in supplementary (Table EV1).

### Development of a T6Es automated XGBoost classifier algorithm

In our efforts to classify potential T6Es proteins effectively, we used a gradient boosting method, XGBoost, which offers an appealing balance between classification accuracy and model interpretability. It is particularly effective on imbalanced datasets where one class, e.g. negative samples, greatly outweighs the other. This capability makes XGBoost particularly well-suited to biological classification tasks, which are often associated with similar challenges (More & Rana, 2017). We used this method before to successfully classify toxins associated with the extracellular contractile injection system (eCIS) (Danov *et al*, 2024a) which shares some characteristics with T6SS, such as phage tail origin.

Our goal was to minimize unnecessary laboratory experiments while still identifying novel T6Es. To achieve this, we optimized our model to maximize Positive Predictive Value (PPV), the proportion of predicted effectors that are true effectors, rather than sensitivity (the ability to capture all true T6Es), thereby reducing false positives. As a result, our optimal model configuration consisted of 100 trees with a maximum depth of 5 and a minimum child weight of 7, corresponding to the minimum number of samples required at each decision node. This configuration achieved a PPV of 0.96, meaning that 96% of the proteins classified as T6Es by the model were true positives. Furthermore, the algorithm demonstrated robust performance, successfully identifying 97% of true effectors in the test set.

We further assessed our model’s performance using a Receiver Operating Characteristic (ROC) curve analysis, which measures how well the model balances true-positive and false-positive predictions. The model achieved an area under the receiver operating characteristic curve (AUC-ROC) of 99%, indicating near-perfect ability to distinguish between T6Es and non-T6Es across different classification thresholds (Fig. 3A).

Furthermore, we computed SHAP values to quantify individual feature contributions to model predictions. Features are ranked on the Y axis by importance, while SHAP values on the X axis represent the magnitude and direction of influence. Positive SHAP values indicate increased probability of T6Es classification, while negative values indicate decreased probability (Fig. 3B). Physicochemical properties emerged as the dominant discriminators of T6E identity. Hydrophobicity ranked as the single most influential feature, with lower values strongly driving effector classification, reflecting the predominantly hydrophilic nature of T6SS effectors. Aromaticity and sequence length followed as the third and fourth most significant contributors, respectively. Mean molecular weight of the residues, charge and structural order further sharpened this picture: T6Es are characterized by lower mean residue molecular weights, indicating a preference for smaller amino acids relative to non-T6Es, alongside a higher proportion of order-promoting residues. Together, these features converge on a coherent physicochemical profile: T6Es tend to be hydrophilic, charged, and enriched in aromatic, low molecular weight, and order-promoting residues.

In addition, domain-based features significantly improved the model accuracy. Several features strongly indicated a non-T6E classification: the presence of core T6SS domains (hcp, VgrG, PAAR) and common bacterial protein domains (the second most important feature). Conversely, certain features supported T6E classification when present: eukaryotic-like domains, glycine zippers (found in pore forming effectors (Ali *et al*, 2023)), and the R COG category (“general function prediction”), although their absence had minimal impact on predictions (Fig. 3B). Genomic location analysis revealed that proximity to T6SS core genes strongly supported T6E classification, whereas greater distances had limited predictive value. These findings indicate that T6Es represent a distinct class of specialized proteins that are genomically co-localized with T6SS machinery and possess unique domain architectures, potentially reflecting their roles as secreted toxins.

Interestingly, several features likely reflects the high variability and poor conservation of immunity genes across strains and species (Hagan *et al*, 2023), with the majority of T6SS immunity proteins remaining uncharacterized to date. Other features did not contribute a lot to the model because they are rare such as having a toxin, MIX (Salomon *et al*, 2014), or FIX (Jana *et al*, 2019) domain annotation (Fig S1G-H).

### New T6Es classification

We used the developed classifier to categorize all 12,726,067 genes extracted from the 2,559 T6SS-encoding bacterial genomes (T6SS+ genomes) (Table EV2). We applied the classifier to this dataset, and each gene received a score ranging from 0 (non-T6Es) to 1 (high-confidence T6Es). Candidate effectors were defined by two criteria: a prediction score exceeding 0.7 and a genomic distance of no more than 8 genes from the nearest core T6SS component (including accessory T6SS loci). Applying these thresholds across the dataset yielded 19,958 putative T6Es (Table EV1)

Functional analysis of the 19,958 predicted T6Es revealed significant structural diversity. While 20.9% lacked identifiable Pfam domains, a subset of 7,526 genes (20.2%) displayed characteristics typical of evolved T6Es, containing VgrG, Hcp, or PAAR domains with N-terminal extensions of at least 100 amino acids. Within this group, the most prevalent was the VgrG domain (PF05954, “Phage GPD”), present in 84.7% of potentially evolved predictions (Sana *et al*, 2015). Additional domain features included RHS repeats (PF05593) in 12.9% of predictions. COG analysis further highlighted this diversity, with 14,475 proteins classified in category R (General function prediction only) and 626 in category S (Function unknown). This remarkable functional heterogeneity among predicted T6Es suggests the existence of novel toxin families, each potentially employing distinct mechanisms of action (Table EV1).

### Three new T6SS effector candidates (T6CEs) kill bacterial and yeast cells intracellularly

To experimentally validate the T6E predictions, we selected eight candidate proteins with high confidence scores (XGBoost probability > 0.70) using stringent filters (see Materials and Methods). Importantly, the selected candidates were “non-trivial” T6SS effector predictions. Namely, they were fairly large proteins (mostly larger than 500 aa) and did not carry signature VgrG/PAAR/Hcp/RHS domains which facilitate manual T6E annotation. The candidates appear in Table 1. We used the fact that most T6Es are accompanied by an unannotated immunity gene which should create a stable protein-protein interaction with the cognate T6E (Geller *et al*, 2024). We prioritized candidates whose AlphaFold3-predicted complexes with neighboring gene products showed medium-to-high interface confidence (ipTM > 0.73; Fig. 4A-C), a metric used to evaluate predicted protein–protein interaction accuracy that can indicate effector–immunity pair (Banhos Danneskiold-Samsøe *et al*, 2024). We named these effector proteins T6ECs standing for type 6 effector candidates.

**Figure 4.**
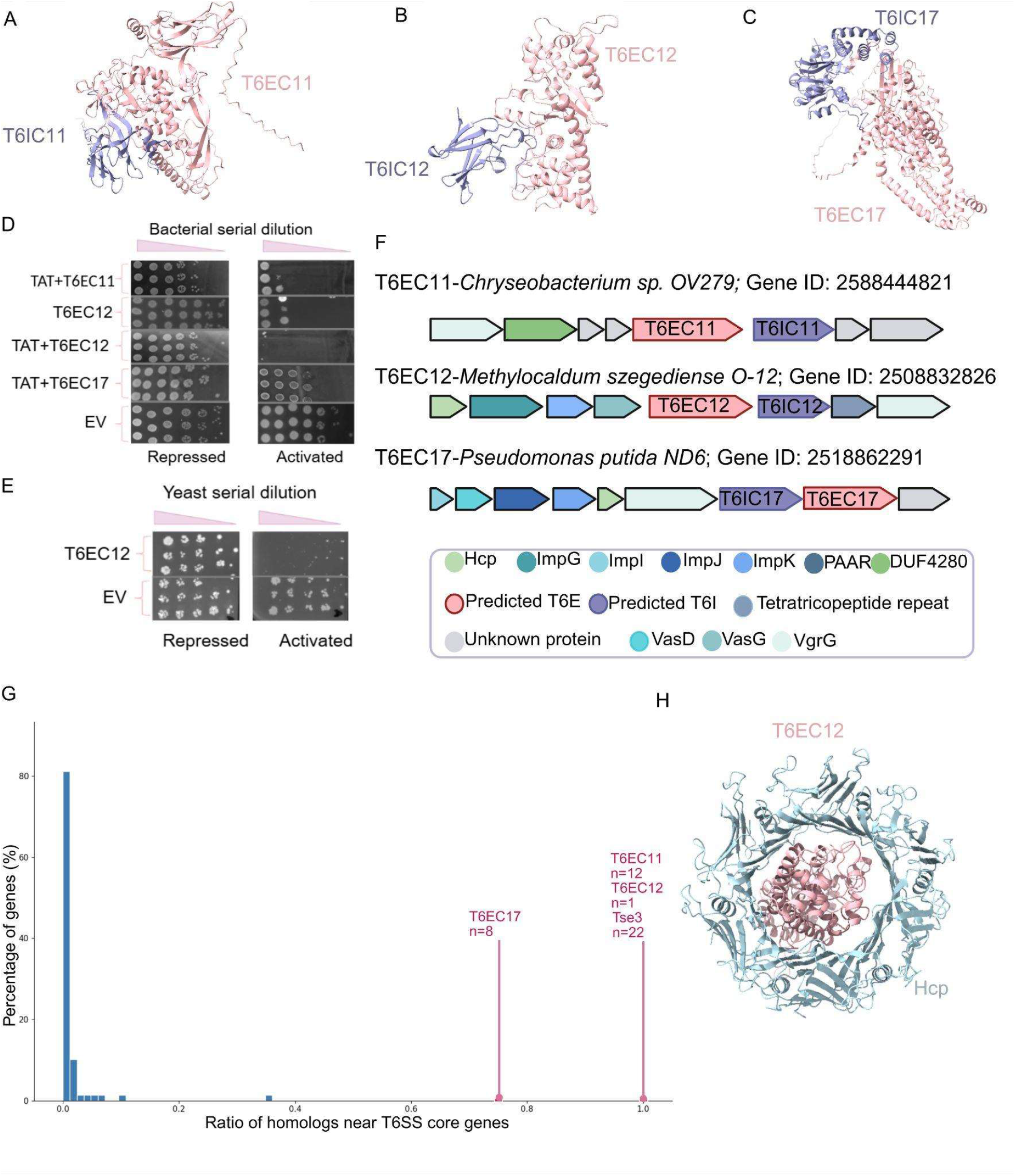
Three novel type VI effector candidates being cytotoxic. **A**. T6EC11 and its predicted immunity T6IC11 with ipTM of 0.81 and pTM=0.76. **B.** T6EC12 and its predicted immunity T6IC12 with ipTM=0.73 and pTM=0.87 **C.** T6EC17 and its predicted immunity T6IC17 with ipTM=0.73 and pTM=0.67. **D.** A representative drop assay of the T6Es that were cytotoxic to *E. coli*. EV, empty-vector as a control. Glucose (1%) and arabinose(0.2%) lead to repression and induction of clonal genes, respectively. Bacteria were serially diluted from left to right to quantify CFUs. **E**. A representative drop assay of yeast T6Es. EV - yeast cell with empty pESC plasmids. Glucose (2%) and galactose(2%) lead to repression and induction of clonal genes, respectively. Yeasts were serially diluted from left to right to quantify CFUs. **F**. Schematic representation and annotation of each validated predicted T6Es and its genomic context. **G**. Ratio of homologs of T6EC11, T6EC12, T6EC17, known T6E Tse3 (pink) and randomly selected genes (blue) from the same genomes located near (within 10 genes upstream or downstream of) at least one T6SS core gene, relative to the total number of homologs identified in our database. **H**. Alphafold3 predicted interaction between T6EC12 and its nearby 6 Hcp with ipTM=0.8 and pTM=0.82.

**Table 1.** List of T6SE candidates.

| Candidate Effector (toxin) Name | ML score | Protein length (aa) | Pfam ID | Genome Name | Genome IMG ID | Gene IMG ID | Confirmed cytotoxicity in *E. coli* cytoplasm | Confirmed cytotoxicity in *E. coli* periplasm | Confirmed cytotoxicity to *S. cerevisiae* |
| --- | --- | --- | --- | --- | --- | --- | --- | --- | --- |
| T6EC11 (“moon toxin”) | 0.80 | 557 | ----- | *Chryseobacterium* sp. OV279 | 2588253712 | 2588444821 | - | + | - |
| T6EC12 | 0.89 | 366 | PF14521 (Aspzincin_M35) | *Methylocaldum szegediense* O-12 | 2508501066 | 2508832826 | + | + | + |
| T6EC13 | 0.87 | 319 | ----- | *Delftia acidovorans* CCUG 15835 | 2541046983 | 2541342974 | - | - | - |
| T6EC14 | 0.93 | 661 | ----- | *Burkholderia silvatlantica* PVA5 | 2510917015 | 2511108423 | - | - | - |
| T6EC15 | 0.70 | 537 | ----- | *Pectobacterium wasabiae* CFBP 3304 | 2537561728 | 2538428094 | - | - | - |
| T6EC16 | 0.92 | 585 | ----- | *Frischella perrara* PEB0191 | 2515154034 | 2515298883 | - | - | - |
| T6EC17 | 0.94 | 688 | PF20249 (MIX) | *Pseudomonas putida* ND6 | 2518645571 | 2518862291 | - | + | - |
| T6EC18 | 0.92 | 563 | ----- | *Elizabethkingia meningoseptica* ATCC 13253 | 2545824725 | 2546582919 | - | Not tested | - |

All eight candidates were tested for toxicity in *E. coli* cytoplasm (Fig. S4-6), seven of which also in the *E. coli* periplasm (Fig. S7-10), and all were tested in *S. cerevisiae* cytoplasm (Fig. S11-12). Not all combinations were tested due to resource constraints. Induction was achieved by adding 0.2% arabinose (*E. coli*) or 2% galactose (*S. cerevisiae*), while repression was achieved by adding 1% glucose (*E. coli*) or 2% glucose (*S. cerevisiae*). Notably, among the eight candidates, three demonstrated significant toxicity against bacteria, yeast, or both, resulting in at least a 100-fold reduction in colony-forming units (CFU, Fig. S4-14). All three effectors, T6EC11, T6EC12, and T6EC17, were located in the vicinity of T6SS core genes (Fig. 4F). To genetically link these toxin genes to the T6SS, we searched for homologs (≥80% identity, ≥80% coverage) and found enrichment near T6SS core genes relative to random genes from the same genomes, as well as relative to the known T6E Tse3 (Hood *et al*, 2010) (Fig. 4G). Using AlphaFold3, we predicted a high-confidence protein–protein interaction (ipTM = 0.8) between T6EC12 and a hexameric Hcp ring, the building block of the T6SS tube, with the effector positioned within the lumen (Fig. 4H), a configuration consistent with previous reports for other T6Es, where the predictions matched experimental validations (Paracuellos *et al*, 2026).

T6EC12 from the moderately thermophilic *Methylocaldum szegediense* O-12 was the most broadly toxic protein in our assays, exhibiting strong toxicity across all tested conditions and causing a ∼10,000-fold reduction in CFU/mL when expressed in both *E. coli* and *S. cerevisiae*. T6EC12 carries an HEXXH zinc-binding motif, characteristic of zinc-dependent metalloproteases, in which the two histidines coordinate the catalytic zinc ion and the glutamate facilitates nucleophilic attack on the scissile peptide bond. Combined with the general toxicity we observed it implies a non-specific proteolytic activity. In contrast, T6EC11 and T6EC17 from *Chryseobacterium sp.* OV279 (isolated from the poplar root rhizosphere) and *Pseudomonas putida* ND6 (isolated from wastewater), respectively, showed toxicity only in the *E. coli* periplasm. When analyzing their homologs, we saw that T6EC11 is mostly found in *Flavobacteria* and *Gammaproteobacteria*. T6EC12 is mostly found in *Betaprotebacteria*, and T6EC17 is found only in *Gammaproteobacteria* (Fig S15-17).

### T6EC11 and T6EC12 expression induce distinct cellular phenotypes in bacteria

To further characterize the cellular effects of T6EC11 and T6EC12 on *E. coli* at the single cell level, toxin-induced morphological changes were examined by fluorescence microscopy at consecutive time points following induction.

Periplasmic localization of T6EC11 resulted in a distinct cell wall-associated phenotype that could be observed when labeling membrane, DNA, and peptidoglycan (Figure 5A). Following induction, cells quickly lost their characteristic rod-shaped morphology and became spherical, a known cell wall damage phenotype already reported by Joshua Lederberg in 1956 (Lederberg, 1956). We expected peptidoglycan degradation to occur relatively uniformly around the cell, resulting in a progressive loss of mechanical stability. Instead, the HADA-labeled peptidoglycan signal became increasingly asymmetric. By 30 min after induction, the remaining peptidoglycan signal was frequently restricted to one side of the cell. Strikingly, in some cells the residual HADA-labeled peptidoglycan was positioned in a large space between separated inner and outer membranes (Fig. 5A, T30, 30 minutes following induction). In some cells, this resulted in a striking spatial organization in which approximately one half of the cell contained DNA whereas the other half contained the residual peptidoglycan. In other cells, the peptidoglycan formed a distinct crescent-shaped structure (Fig. 5A, T30).

**Figure 5.**
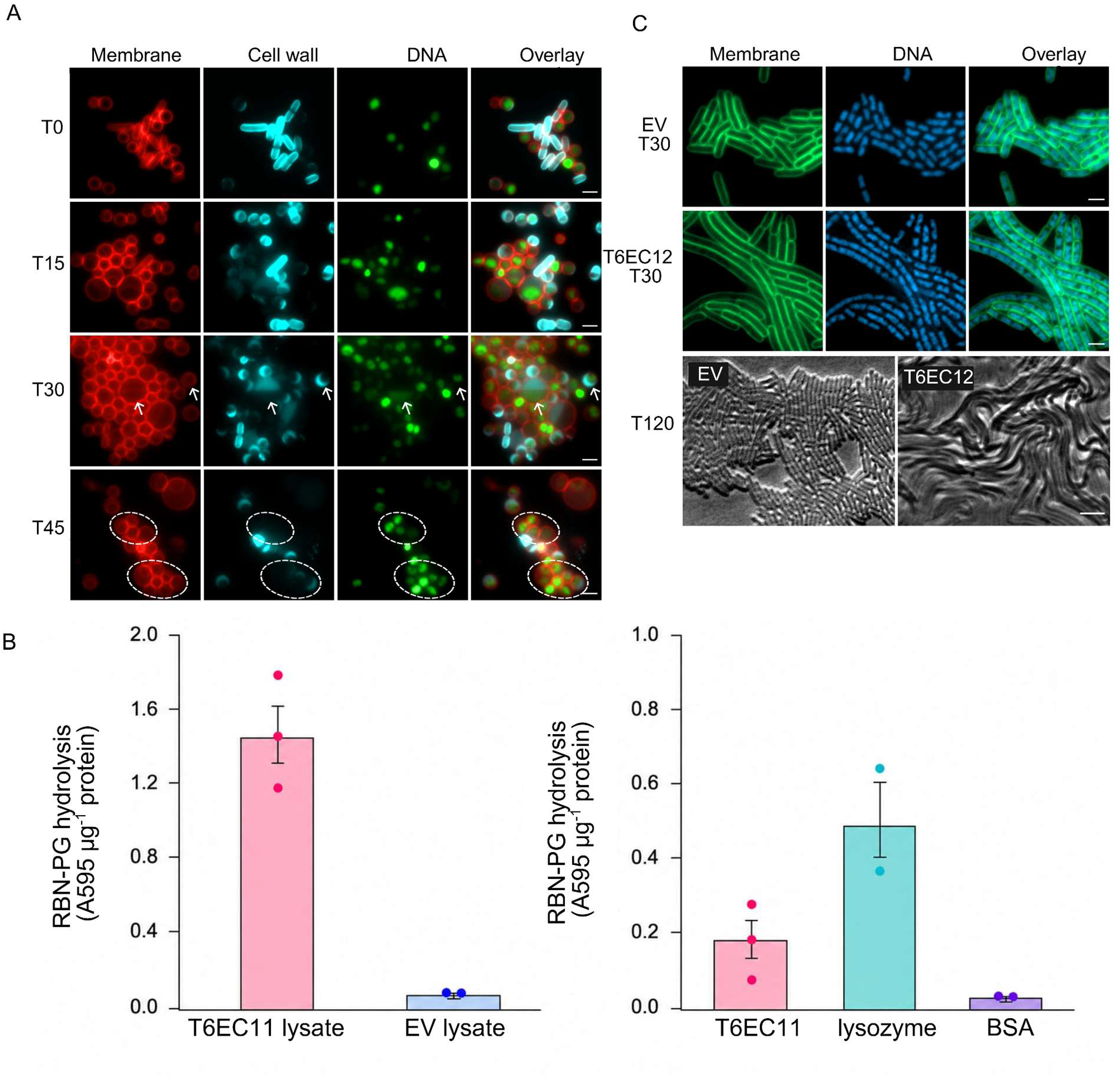
T6EC11 and T6EC12 expressions induce distinct cellular phenotypes in *E. coli,* and T6EC11 degrades peptidoglycan. **A.** T6EC11 induces asymmetric peptidoglycan loss and a moon-like phenotype in *E. coli*. T6EC11 was expressed in the periplasm of *E. coli* BL21(DE3), and cells were visualized by fluorescence microscopy at the indicated times following induction with 0.1% arabinose. Peptidoglycan was labeled with HADA (cyan) prior to toxin induction, while the cytoplasmic membrane was stained with FM4-64 (red) and DNA with SYTO9 (green). Arrows indicate moon-like cells. Dashed circles indicate cells containing DNA and membrane but lacking peptidoglycan. Images are representative of five independent replicates. Scale bar, 2 μm. Txx - xx minutes following induction. **B.** Peptidoglycan hydrolytic activity of T6EC11. Peptidoglycan hydrolysis by purified T6EC11 and lysates of T6EC11 expressing *E. coli* was assessed using RBB-labeled peptidoglycan. Hydrolytic activity was quantified by measuring the absorbance of soluble RBB-containing fragments at 595 nm. EV lysate represents *E. coli* harboring the empty vector, serving as the corresponding lysate control. Lysozyme and BSA serve as controls for purified T6EC11. Values were normalized to the amount of protein added to each reaction and are expressed as A595 µg⁻¹ protein. Bars represent mean ± SD, and dots represent individual replicates. For purified T6EC11 and T6EC11 lysate, n = 4; for EV lysate, lysozyme, and BSA, n = 2. **C.** T6EC12 leads to filamentous bacteria formation. T6EC12 was cloned and grown as described for T6EC11. Overlay images of DNA (blue) and membrane (green) stained in *E.coli* cells are shown. Empty-vector control cells exhibited normal growth and cell division (EV T30 and EV T120), whereas T6EC12-expressing cells displayed pronounced elongation without dividing following induction (T6EC12 T30 and T6EC12 T120). Txx - minutes following induction. Representative images from a single replicate out of three independent replicates are shown. Scale bar: 2 μm.

As the phenotype progressed, the detectable peptidoglycan signal was almost completely lost in a subset of cells by 45 min after induction (Fig. 5A, T45). Remarkably, DNA and membrane signals remained detectable at this stage, indicating that loss of the peptidoglycan signal preceded complete disruption of cellular organization. Thus, T6EC11 causes a highly asymmetric and progressive loss of cell-wall-associated peptidoglycan rather than the uniform degradation initially anticipated. This magnificent polar cell wall gradual disappearance convinced us to name T6EC11 the “moon toxin” due to its resulting crescent-like bacterial phenotype.

The progressive loss of peptidoglycan observed by microscopy prompted us to test whether T6EC11 directly possesses peptidoglycan-hydrolytic activity. To address this, we measured the release of Remazol Brilliant Blue (RBB) from RBB-labeled peptidoglycan. Both purified T6EC11 and lysates from T6EC11 expressing *E. coli* exhibited increased peptidoglycan hydrolysis relative to their respective controls (Fig. 5B). Namely, purified T6EC11 showed higher hydrolytic activity than BSA, while T6EC11 expressing cell lysates showed ∼150 fold higher peptidoglycan degradation activity than the empty vector (EV) lysate. Lysozyme, included as a positive control, also showed peptidoglycan hydrolysis. Together, these results support T6EC11 as a novel peptidoglycan hydrolase with enzymatic activity toward bacterial peptidoglycan.

In parallel, T6EC12 expression resulted in progressive cell elongation and filamentation (Fig. 5C). At 30 min after induction, T6EC12-expressing cells were already markedly elongated. Membrane staining showed continuous elongated cell boundaries, while DNA staining revealed multiple nucleoids distributed along the length of the filaments. By 120 min, the phenotype became substantially more pronounced, with extensive, highly elongated filamentous cells observed throughout the population. These observations indicate that T6EC12 expression progressively inhibits cell division while allowing continued cellular elongation, consistent with disruption of the bacterial division or septation process.

### T6EC12 contains a conserved HExxH catalytic center within a novel two-domain scaffold

To investigate the molecular mechanism underlying T6EC12 toxicity, we first examined the toxin’s predicted catalytic region. Sequence analysis identified a conserved HExxH motif (H301-E302-L303-S304-H305), a hallmark of zinc-dependent metalloproteases, together with two highly conserved acidic residues (D312 and E341) located immediately downstream of the motif (Fig. 6A). Examination of the AlphaFold model revealed that these residues converge within a prominent solvent-accessible cleft, forming a well-defined putative active site pocket (Fig. 6B, and S19A). Surface analysis demonstrated that this region forms a deep substrate-accessible channel lined by the predicted catalytic residues and characterized by a predominantly negatively charged electrostatic potential (Fig. S19A). Adjacent to the catalytic pocket, a cluster of aromatic and hydrophobic residues form a hydrophobic cavity that may contribute to substrate recognition and specificity (Fig. 6B). Notably, the predicted immunity protein, T6IC12, binds directly over this region, effectively occluding the catalytic cleft and preventing access to the putative active site (Fig. S19B), consistent with the common mechanism in which immunity is achieved through steric inhibition of catalysis.

**Figure 6.**
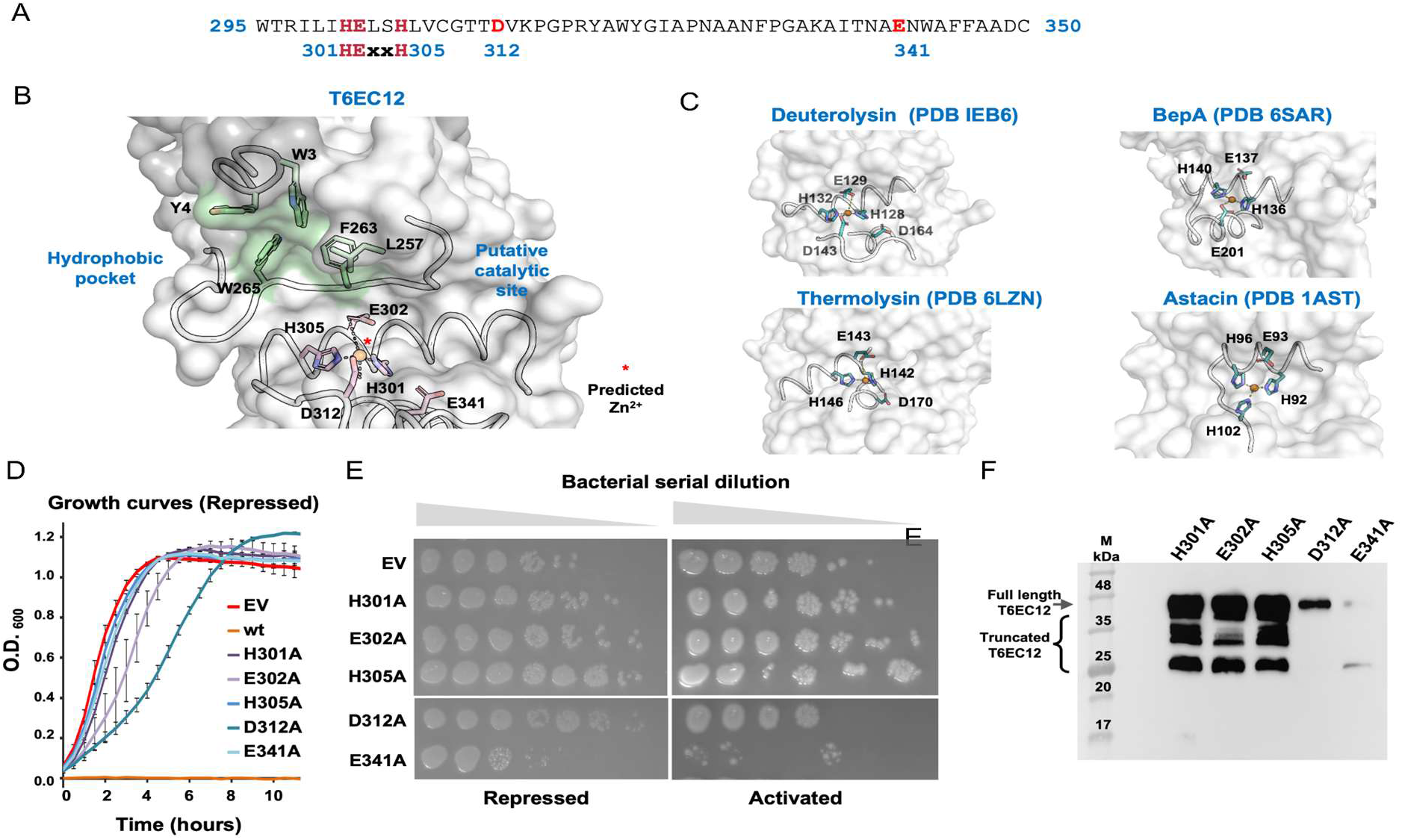
Structural and functional characterization of the predicted catalytic center of T6EC12. **A.** Amino acid sequence surrounding the predicted catalytic center of T6EC12. The conserved HExxH motif (H301-E302-L303-S304-H305) and the neighboring acidic residues D312 and E341 are highlighted. **B.** AlphaFold model of the T6EC12 catalytic region showing the putative active-site pocket. The predicted catalytic residues are shown as sticks and colored in pink, and the adjacent hydrophobic pocket is highlighted and colored in green. **C.** Structural comparison of the predicted T6EC12 active site with representative HExxH metalloproteases. Comparison of the catalytic centers of T6EC12, Deuterolysin, Thermolysin, BepA, and Astacin (see Fig. S18C) demonstrates conservation of the HExxH-centered active-site geometry despite differences in the overall protein architecture. **D**. Growth curves of *E. coli* BL21(DE3) cells expressing wild-type T6EC12 or catalytic-site variants in repressed condition. wt T6EC12 abolished bacterial growth, whereas some catalytic-site mutants exhibited delayed growth relative to the empty-vector control. **E.** Drop assay of *E. coli* BL21(DE3) expressing wild-type T6EC12 or catalytic-site variants under repressing (2% glucose) and inducing (0.01 mM IPTG) conditions. Catalytic-site mutants retained the ability to grow upon induction. **F.** Immunoblot analysis of catalytic-site variants following small-scale expression. Multiple discrete protein species were detected in addition to the full-length protein.

To assess whether the predicted T6EC12 active site resembles those of known metalloproteases, we compared its AlphaFold model with representative HExxH-containing proteases for which experimental structures are available. Although the overall architecture of T6EC12 did not closely resemble that of the examined metalloprotease families, superposition of the active-site regions revealed conservation of the HExxH-centered catalytic geometry, including the predicted zinc-coordinating histidines and adjacent acidic residues (Fig. 6C). In contrast to the compact single-domain architecture of several characterized HExxH metalloproteases, T6EC12 is predicted to comprise two distinct domains separated by an interdomain region (Fig. S19A), with the putative catalytic pocket located in the C-terminal domain and positioned near the domain’s interface. These observations suggest that T6EC12 adopts a conserved HExxH metalloprotease-like active-site architecture within a previously uncharacterized two-domain structural scaffold.

### Experimental validation of the predicted catalytic center of T6EC12

To experimentally evaluate the functional importance of the predicted catalytic center, the T6EC12 gene was cloned into the pET28 expression vector. Alanine substitutions were introduced into the HExxH motif residues H301, E302, and H305, and the neighboring acidic residues D312 and E341. Repeated attempts to transform the wild-type (wt) T6EC12 construct into *E. coli* BL21(DE3) cells resulted with no or only few colonies, even under suppression conditions (2% glucose), indicating that low-level expression of the wt toxin is highly detrimental to the host. In contrast, the T6EC12 mutants yielded transformants, although transformation efficiency remained reduced compared to an empty vector.

Growth experiments in liquid media indicated that some of the T6EC12 mutants delayed bacterial growth whereas the wt abolished growth by its unclonability (Fig. 6D). To assess the contribution of the predicted catalytic residues to T6EC12 toxicity, a drop assay was performed under inducing conditions. All catalytic-site mutants retained the ability to grow upon induction, although the extent of growth varied among the variants (Fig. 6E), indicating that residues forming the predicted HExxH-centered active site are required for the full toxic activity of T6EC12. The catalytic-site variants were subsequently subjected to small-scale expression followed by immunoblot analysis. The H301A, E302A, and H305a mutants were readily expressed, with lower expression of the D312A and E341 mutants. However, immunoblotting consistently revealed multiple discrete truncated protein species in addition to the full-length protein (Fig. 6F), suggestive of auto-proteolysis.

## Discussion

Research into T6Es is crucial for understanding bacterial competition and virulence, and developing biotechnological applications, such as new antimicrobial strategies, genome editing tools (Mok *et al*, 2020), enhanced colonization of probiotics, and pest control methods. However, identifying T6Es is challenging due to their diverse mechanisms and the absence of a characteristic signal peptide. In addition, most works focus on the T6Es of human pathogens although T6SS is prevalent across gram-negative bacteria. To classify new T6Es, we have developed an XGBoost model that improves upon previous sequence-based methods by incorporating genomic context, biochemical traits, and protein function annotations.

To date ML models for predicting T6Es were focused on what can be understood from the protein sequence alone (Sen *et al*, 2019; Jiaweiwang *et al*, 2018; Hu *et al*, 2025), although almost all identified effectors are encoded with genetic linkage to the T6SS main gene cluster(s) or orphan VgrG/PAAR islands (Lin *et al*, 2013; English *et al*, 2012; Abby *et al*, 2016; Carobbi *et al*, 2024). Our model demonstrates that the location of the gene is of high importance in its classification as T6E (Fig. 1A).

Our analysis revealed that biochemical traits are highly influential in T6E classification, indicating that these T6Es possess distinct physicochemical properties. They tend to be hydrophilic, charged, and enriched in order-promoting and aromatic amino acids, with a preference for lighter amino acids (Fig. 2). This physicochemical signature was yet to be discussed and can help with the characterization of new effectors. T6Es are also strongly associated with certain protein COG categories: R (General Function Prediction Only) and X (Mobilome, likely to VgrG and PAAR extensions) (Fig. S1).

One feature that worked surprisingly well is “Common Domain Presence” (Figs. 1B, 3B) that helped us to reduce false positives because it is strongly anticorrelated with T6Es. Given that most random genes within T6SS+ genomes are non-T6Es due to T6E low abundance in a given genome, and the fact that T6Es often carry unique enzymatic functions, it made sense to label highly abundant genes as non-T6E genes. We believe that this is a useful method to employ in other machine learning classification tasks in which one tries to classify a “needle in a haystack”: characterize the common appearance of hay.

Using our model, we predicted 19,958 putative T6Es. To enrich for high-confidence candidates, we filtered results by prediction score (>0.7) and genomic proximity (≤8 genes from the nearest core T6SS component), then tested eight candidates for their toxicity to *E. coli* or *S. cerevisiae* to assess model accuracy. To identify new T6E classes we looked at T6E candidates from poorly studied environmental microbes (Table 1), that are relatively long and lack similarity to known T6Es. We also avoided focusing on evolved T6Es which are easy to identify.

The “moon toxin” (T6EC11) showed strong toxicity when expressed in the periplasm (Fig. 4D, Fig S7) but not in the cytoplasm (Fig. S4), with only modest rescue by the predicted immunity protein (Fig S13). To gain insight into its possible mode of action, we examined *E. coli* cell morphology in the presence and absence of toxin expression. Cells expressing the moon toxin in the periplasm displayed rounding and apparent loss of peptidoglycan integrity, suggesting that T6EC11 may degrade peptidoglycan (Fig. 5A), an enzymatic activity we indeed confirmed (Fig. 5B). To the best of our knowledge, we present the first indication that peptidoglycan is degraded in a polar gradual manner portrayed as a crescent-like phenotype. We propose that the moon toxin accumulates in one pole and either pushes or pulls the peptidoglycan as part of its rapid degradation. Interestingly, DNA and the membrane remain stable while the peptidoglycan in the cell is fully degraded. It would be interesting to study whether other peptidoglycan hydrolases and amidases are acting in the same polar fashion within the periplasm.

Compared with the abundant nuclease, phospholipase and cell wall-targeting T6SS effectors described to date, protease-like effectors remain relatively uncommon. The identification of T6EC12 therefore expands the known catalytic diversity of T6SS toxins. T6EC12 showed high toxicity in every assay tested, namely killing of bacteria (cytoplasm and periplasm) and eukaryotic cells. Although the Aspzincin_M35 domain (PF14521)(Rocha *et al*, 2026), found in T6EC12 is an established toxic domain, T6EC12 diverges substantially in sequence and predicted structure from previously characterized members, and its mechanism of action remains unresolved. A homolog sharing only 30% sequence identity, with a totally different immunity protein, was recently identified among antibacterial T6Es from *P. aeruginosa* and *E. coli*(Wang *et al*, 2023) (Fig S18), suggesting that members of this family have evolved multiple, non-conserved mechanisms of toxin neutralization. Consistent with this notion, structural modeling predicts that the T6EC12 immunity protein sterically occludes the putative active-site cleft (Fig S19B), suggesting inhibition through direct blockage of substrate access. Members of the M35 family have also been reported to exhibit toxicity toward yeast and insects through a proposed protease-like mechanism, although their molecular targets and enzymatic activities remain largely unresolved (Huang *et al*, 2020).

We were unable to directly demonstrate proteolytic activity of T6EC12 due its high toxicity that prevented purification of the full length protein. However, alanine substitutions within the predicted HExxH catalytic center and its neighboring residues consistently reduced toxin-mediated growth inhibition. Together with the structural conservation of the active-site geometry, these findings support the functional importance of the predicted catalytic center and are consistent with the hypothesis that T6EC12 belongs to a previously unrecognized HExxH metalloprotease-like family of T6SS effectors.

Expression of T6EC12 resulted in progressive cell elongation and inhibition of cell division (Fig. 5C). Because filamentation is a common bacterial response to diverse cellular stresses (Gates, 1933; Ting *et al*, 2018; Sánchez-Gorostiaga *et al*, 2016), this phenotype alone does not identify the molecular target of T6EC12. Elucidating the physiological substrate and biochemical activity of this toxin will therefore require further investigation.

While our classification tool achieved high predictive accuracy, false negatives and positives remain possible, given the classifier’s reliance on the quality of the training data and the relatively limited positive training set. Heterologous expression experiments confirmed cytotoxic activity for 37.5% of selected candidates, lower than the model’s reported precision (96%), though based on a small sample size. The five candidates lacking cytotoxicity may represent bona fide secreted effectors with non-toxic functions, as previously documented for some T6Es (Si *et al*, 2017; Lin *et al*, 2017; Zhu *et al*, 2021). Alternatively, assay sensitivity may have been limited by testing in suboptimal target organisms, cellular compartments, or conditional toxicity (LaCourse *et al*, 2018). For example, TseH, a cysteine endopeptidase, was shown to be toxic to *Aeromonas* species but not to *E. coli* due to presence of envelope stress-response pathways in *E. coli* (Hersch *et al*, 2020b). Further validation across diverse organisms and cell types is warranted to improve detection and elucidate target specificity, but is beyond the scope of this work.

In conclusion, the successful identification and validation of novel T6Es using a newly developed machine learning approach expands our understanding of T6SS functional diversity. The resource of over 19,000 predicted T6Es provides numerous hypotheses for future investigation, with potential to advance fundamental microbiology.

## Materials and Methods

All samples and their corresponding features were obtained from the Integrated Microbial Genomes and Microbiomes database (IMG) (Markowitz *et al*, 2012). All the code available in https://github.com/AvitalAkermanArad/T6SS_ML_Project

### 1. Data collection, preprocessing, and compilation of T6Es and non-T6Es sets

#### i) Assembling Positive Datasets (T6Es)

We compiled the positive set using data from the Secret6 database of T6Es (Li *et al*, 2015), which includes 335 validated entries with 74,712 bioinformatically predicted T6Es. Subsequently, to retrieve the T6Es genes and proteins, we identified homologous sequences with a 100% similarity for these entries from the IMG database by employing Diamond (Buchfink *et al*, 2021) for protein similarity. Features were then extracted based on the information available in the IMG database. To eliminate redundancy in the sequence data and ensure high data quality, we excluded metagenomes and highly homologous genes (with identity above 80%) by employing cd-hit (Fu *et al*, 2012) to ensure sequence diversity within the resulting positive set. The final positive set consisted of 634 genes, including 79 validated effectors and 555 putative effectors. Note the large fraction of putative effectors.

#### II) Assembling Negative Datasets (non-T6Es)

Our negative set consisted of three different categories: (1) 377 non-homologous sequences of effectors of type 1 to 4 secretion systems (Wang *et al*, 2021). (2) 672 core T6SS genes, defined as encoding structural components of the secretion system, but not evolved effectors. (3) 2,939 randomly selected non-toxin genes from genomes encoding type VI secretion systems ("T6SS+"). Given the low prevalence of T6Es in microbial genomes relative to non-T6Es, the probability of inadvertently including unidentified T6Es in these random selections was deemed negligible. This random sampling approach was implemented to expose the model to diverse genetic variations.

### 2. Feature extraction

For each protein, a comprehensive set of features was computed, encompassing genome-based and protein sequence-based attributes (Table EV1):

#### Biochemical features

were derived directly from the protein sequence and included protein length, along with several length-normalized properties calculated by summing per-residue values across the sequence and dividing by sequence length. Mean hydrophobicity was calculated using the Kyte-Doolittle hydrophobicity scale (Kyte & Doolittle, 1982), which assigns positive values to hydrophobic residues. Mean charge was calculated by assigning +1 to positively charged residues (R, H, K), −1 to negatively charged residues (D, E), and 0 to all others. Mean disorder/order-promoting index was calculated by assigning +1 to disorder-promoting residues (S, A, Q, E, R, K, G, P) and −1 to order-promoting residues (I, L, M, V, F, Y, C), based on their established tendency to hinder or facilitate protein folding (Campen *et al*, 2008). Mean molecular weight was calculated by summing individual residue molecular weights, and mean aromaticity was calculated as the proportion of aromatic residues in the sequence.

#### Genomic features

were derived from domain composition, genomic context, and annotation databases, with most encoded as binary indicators. Toxin domain presence was assessed using HMMER3 (Eddy, 2009) against the Toxinome database (Danov *et al*, 2024b), assigning a value of 1 if any toxin domain was detected. Type VI core domain presence was evaluated against 35 core T6SS Pfam domains (Mistry *et al*, 2021)(Supp. Table S1). to exclude proteins resembling core structural components. A "potentially evolved T6SE" feature flagged proteins likely to have arisen through domain fusion (Geller *et al*, 2021), assigned a value of 1 when T6SS structural domains (Hcp, PAAR, or VgrG) were present at the N-terminus and at least 100 additional C-terminal residues remained available to accommodate an unannotated toxin domain. Eukaryotic-like domain (ELD) presence was assessed using domains curated from EffectiveDB (Eichinger *et al*, 2016) and validated via InterPro (Paysan-Lafosse *et al*, 2023), with domains classified as eukaryotic-like if their prevalence in eukaryotic organisms exceeded 50% (Table EV1), reflecting their potential role in host-cell interaction through molecular mimicry (Mondino *et al*, 2020).Glycine zipper presence was scored based on the occurrence of the GXXXG motif within the first 50 N-terminal residues, a feature associated with membrane-active T6Es and pore-forming toxins (Ali *et al*, 2023). A common domain presence feature, hypothesized to be anticorrelated with effector identity, was defined using Pfam domains extracted from ∼12 million genes across T6SS+ genomes in the IMG database; proteins containing one of the 25% most frequent domains were assigned a value of 1, reflecting the reduced likelihood that widely distributed domains represent specialized effector functions(Table EV1). Mobile element domain presence was similarly scored to filter out non-core, transposon-associated false positives(Table EV1). MIX and FIX domain presence was determined by RPS-BLAST against the NCBI Conserved Domain Database (Yang *et al*, 2020), screening for five characterized domain variants using stringent thresholds (E-value < 0.001, sequence identity > 40%). Signal peptide presence was obtained from IMG annotations, as its presence indicates Sec-dependent secretion rather than T6SS-mediated export, given that T6Es lack canonical signal sequences (Lewis *et al*, 2019; Liang *et al*, 2015).COG category presence was assigned using the standard 26-category COG classification system (Galperin *et al*, 2021), with differential representation between positive and negative sets evaluated by chi-square test (Tallarida & Murray, 1987) with FDR correction(Benjamini & Hochberg, 1995).Distance from the nearest T6SS core Pfam (Table S2) was calculated as the number of intervening genes, reflecting the frequent genomic co-localization of T6Es with core T6SS components(Alcoforado Diniz *et al*, 2015).proteins lacking a nearby core gene were assigned a value of −1. Finally, immunity domain presence was assessed in the immediately flanking genes (one upstream, one downstream), with a value of 1 assigned if either contained a known immunity domain (Danov *et al*, 2024b), consistent with the conserved genetic co-localization of T6SS effector-immunity gene pairs (Dong *et al*, 2013).

### 3. Feature selection

COG category U (intracellular trafficking, secretion, and vesicular transport) was excluded from the final model despite significant discriminatory power (Fig. S1), due to dataset composition bias: this annotation was 1.5-fold overrepresented in the negative dataset (T6SS core components and other secretion system effectors) and, within the positive dataset, was largely restricted to evolved T6Es. Retaining this feature risked biasing the model against novel T6Es, particularly evolved effectors and cargo proteins, based on dataset artifacts rather than biological signals.

### 4. Model selection

Following our successful previous experience in effector classification(Danov *et al*, 2024a), we selected the XGBoost (eXtreme Gradient Boosting) algorithm(Chen & Guestrin) for our classification task. XGBoost works like building a strong team of simple "expert" decision trees, where each new expert focuses on correcting the mistakes of the previous ones. Hyperparameter tuning was performed to optimize the algorithm’s performance and minimize false positives. Optimized hyperparameters include the number of trees per classifier (n_estimators=100), the maximum tree depth (max_depth=5), and the minimum child weight (min_child_weight=7). We used the XGBoost Python package, with optimal hyperparameters.

### 5. Performance evaluation

Model reliability was assessed using fivefold cross-validation, with the dataset split into five stratified segments preserving the T6E-to-non-T6E ratio; in each iteration, 80% of the data was used for training and 20% for validation, rotating across all segments. Performance was evaluated using positive predictive value (PPV, precision) and sensitivity. Since our primary goal was to optimize resource allocation for downstream experimental validation, PPV was prioritized over sensitivity to maximize the success rate of validation experiments.

To independently confirm generalizability, the model was tested on a held-out set of 231 proteins (5% of each class, randomly selected, preserving the original effector-to-non-effector ratio) not used in training or cross-validation. The model achieved a PPV of 96% on this independent set, consistent with training performance, indicating that it learned generalizable patterns rather than overfitting to the training data.

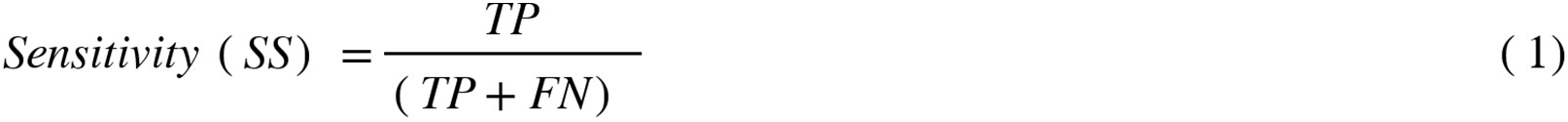

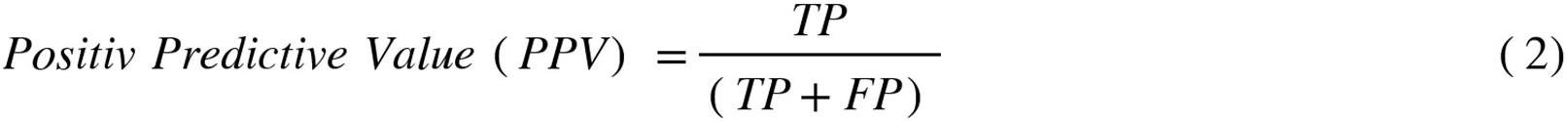

### 6. New T6Es classification

The trained model was applied to all genes derived from genomes that contain T6SS (T6SS+ genomes)(Geller *et al*, 2021). Features of these genes were extracted from IMG, and the trained model was employed to predict their classification as T6E (1) or non-T6E (0) based on the calculated score, with a score > 0.70 indicating a T6E prediction. To reduce false positives, only genes located within a genomic window of ±8 genes relative to any known T6SS-associated gene were retained as putative T6Es..

### 7. Selection of candidates for experimental validation

The selection of candidates for experimental validation was based on a systematic filtering approach. Based on the predictions of our high-confidence XGBoost model (scores > 0.70), we excluded potentially evolved proteins and those with ≥ 50% sequence identity to our positive training set. The remaining candidates were clustered using MMseqs2 (Steinegger & Söding, 2017) at 50% sequence identity and evaluated based on multiple criteria: Proximity to T6SS core genes, presence of neighboring genes with potential antitoxin function (identified by AlphaFold 3(Abramson *et al*, 2024) protein-protein interaction predictions), protein length (prioritizing shorter sequences to reduce DNA synthesis costs), and domain architecture. Most selected candidates lacked known domains and showed no structural similarity to known proteins using Foldseek (van Kempen *et al*, 2023). Ultimately, we identified eight candidates for experimental validation in bacterial and yeast toxicity assays.

### 8. Experimental validation

All primers used in this project are in Table S3.

#### Bacterial strains and strain construction

Candidate T6SE gene sequences were retrieved from IMG, synthesized with codon optimized for expression in *E. coli*, and cloned into pBAD24 by Twist Bioscience (South San Francisco). The growth control for the experiments was an empty-vector. The plasmid was then transformed into *E. coli* BL21 (DE3) strain, using the previously described TSS method(Chung *et al*, 1989). For the toxin-immunity protection assay, *E. coli* BL21(DE3) cells harboring pBAD24 plasmids with toxic genes were co-transformed with pET28 vectors harboring cognate immunity genes or with empty-vector as a control.

#### Assessing antibacterial toxicity in the E. coli cytoplasm

Bacterial drop assays were conducted using the pBAD or the pET28 vectors. For drop assay using the pBAD vector, overnight cultures were grown in LB supplemented with with 1% glucose and 100 µg/ml ampicillin with shaking (250 rpm). The next day, the culture was diluted 1:100 into fresh medium supplemented with the appropriate antibiotic and 0.1% glucose, incubated at 37 °C until OD₆₀₀ of 0.4, and serially diluted by a factor of ten. Dilutions were spotted as three biological replicates on LB plates with ampicillin and 0.2% arabinose as an inducer, or with 1% glucose as a repressor, for cells containing the toxin candidates. Drop assay using the pET28 vector were conducted in a similar protocol with 50 µg/ml kanamycin for selection, 0.01mM IPTG as an inducer and 2% glucose as repressor. The plates were incubated overnight at 37 °C. Results were documented using Amersham ImageQuant 800.

#### Assessing antibacterial toxicity in the *E. coli* periplasm

To test the action of a toxin within the E. coli periplasm, a twin-arginine translocation (TAT) sequence was inserted into the plasmid at the N-terminus to the toxins T6EC11-T6EC17:

MNNNDLFQASRRRFLAQLGGLTVAGMLGPSLLTPRRATAAQA

The rest of the drop assay was performed as described in “Assessing antibacterial toxicity in the *E. coli* cytoplasm”

#### Assessing anti-eukaryotic toxicity to yeast cells

Gene sequences of the toxins that we suspected might be anti-eukaryotes were tested for toxicity to yeast. The genes were retrieved from IMG, amplified by colony PCR and ligated into a cut pESC-leu (with XhoI) plasmid using NEBuilder. The plasmids were then transformed into *Saccharomyces cerevisiae* BY4742 strain. Overnight cultures of the strains containing the vectors of interest were grown in SD-Leu media. For heterologous expression, the culture OD was normalized to OD₆₀₀ of 0.3, washed once with PBS, and then serially diluted by a factor of ten. Dilutions were spotted as three biological replicates on SD-leu agar containing 2% glucose as a repressor or 2% galactose as an inducer, and plates were incubated for 72 h at 30 °C. Results were documented using Amersham ImageQuant 800.

#### Rescue assay by immunity genes

T6IC11 and T6IC12 were cloned into pET28 by Twist Bioscience (South San Francisco). Using heat shock, pET28 + immunity genes were transformed into *Escherichia coli* BL21. Transformed colonies were then used to create competent cells, which were in turn transformed with pBAD24 + toxin genes (T6EC11 and T6EC12). Overnight cultures of the strains harboring the vectors of interest were grown in LB containing kanamycin and ampicillin. Cultures were normalized to OD₆₀₀ of 0.4 and subsequently serially ten-fold diluted. Dilutions were spotted on LB agar containing the proper selection and inducer; plates with either 1% Glucose and 0.01 mM IPTG, or 0.2% Arabinose and 0.01 mM IPTG.

#### Fluorescence microscopy

Microscopy was performed on induced *E. coli* BL21 cells containing T6EC11 and T6EC12 grown in LB media supplemented with the appropriate antibiotic at 37 °C overnight. Overnight cultures were diluted 1:100 into fresh LB medium and grown for 2 h until reaching the log phase. For T6EC11 strain fluorescent d-amino acid 7-hydroxycoumarincarbonylamino-d-alanine (HADA) (Tocris Bioscience) was then added to the cultures at a final concentration of 500 μM for 1 hour to allow incorporation of HADA into newly synthesized peptidoglycan as previously described(Mamou *et al*, 2022) with minor modifications. Following incubation, cells were centrifuged at 8,000×rpm, and the pellet washed with PBS (x3) to remove excess unincorporated HADA. The cell pellet was resuspended in 70 µl of PBS with 7 μg/mL membrane stain FM4-64 (Thermo Fisher Scientific T13320) and 0.7 μM nucleic acid stain SYTO 9 (Thermo Fisher Scientific S34854). Cells were then transferred to a chamber slide (VWR Scientific) filled with M9 medium containing 0.1% arabinose,, 0.875 μg/mL membrane stain FM4-64, 2mM *MgSO_4_*, 0.1mM *CaCl_2_*, 1mg/ml *NH_4_Cl*, 0.05% Casamino acids, 1% agarose pads.

For T6EC12 strain, bacteria were induced for 40 min with 0.2% arabinose. Next, 200 µL of cells were centrifuged at 10,000×g, and the pellet was resuspended in 50 µl of PBS with 1 μg/ml membrane stain FM1-43 (Thermo Fisher Scientific T35356) and 2 μg/ml DNA stain 4,6-diamidino2-phenylindole (DAPI; Sigma-Aldrich D9542-5MG).

Cells were visualized and photographed using an Axioplan 2 microscope (Zeiss) equipped with an ORCA-Flash4.0 camera (Hamamatsu). System control and image processing were carried out using ZEN software version 3.1 (Zeiss).

#### RBB-labeled peptidoglycan hydrolysis assay

##### Protein expression and cell lysate preparation

Overnight cultures of *E. coli* BL21 expressing T6EC11 from the pBAD24 plasmid were diluted 1:100 into 100 mL LB medium and grown at 37°C with shaking at 250 rpm for OD₆₀₀ of 0.5. Protein expression was induced by the addition of L-arabinose to a final concentration of 0.02%, and cultures were incubated for an additional 3 h at 37°C with shaking at 250 rpm. Cells were harvested by centrifugation at 4,000 rpm for 20 min at 4°C and maintained on ice throughout lysate preparation. Cell pellets were resuspended in a 10 mL lysis buffer containing 50 mM Tris-HCl (pH 7.4) and 150 mM NaCl. The cell suspension was transferred to 2-mL FastPrep Lysing Matrix B tubes (MP Biomedicals, Cat. No. 116911100) and disrupted using a FastPrep bead beater for two 40-s cycles at 6 m s⁻¹. Cell debris was removed by centrifugation at 15,000 × *g* for 15 min, and the cleared supernatant was collected.

##### FLAG affinity purification

FLAG-tagged T6EC11 was purified from cleared cell lysates using Anti-FLAG M2 Magnetic Beads (Sigma-Aldrich, Cat. No. M8823). Beads were equilibrated by washing twice with lysis buffer and incubated with the cleared lysate for 2 h at 4°C with gentle end-over-end rotation. The beads were collected using a magnetic rack, and the unbound fraction was removed. Protein-bound beads were washed four times with a lysis buffer. FLAG-tagged T6EC11 was eluted by incubation with 150 ng µL⁻¹ 3×FLAG peptide in lysis buffer for 30 min at room temperature. The beads were magnetically separated, and the supernatant containing the purified protein was collected. Protein concentrations of purified fractions and cell lysates were determined using Bradford Reagent (Sigma-Aldrich, Cat. No. B6916) according to the manufacturer’s instructions. Samples were diluted as required to fall within the linear range of the assay.

##### RBB-PG hydrolysis assay

Peptidoglycan from *Bacillus subtilis* (Sigma-Aldrich, Cat. No. 69554) was covalently labeled with 20 mM Remazol Brilliant Blue (RBB; Sigma-Aldrich, Cat. No. R8001) in 0.25 M NaOH at 37°C overnight. The reaction was neutralized with 0.2 M HCl, and RBB-labeled peptidoglycan was collected by centrifugation at 20,000 × *g* for 15 min at room temperature. The pellet was repeatedly washed with Milli-Q water until no soluble RBB was detected in the supernatant following centrifugation. For peptidoglycan hydrolysis assays, 20 µL of RBB-labeled peptidoglycan was incubated with the indicated purified protein or cell lysate in reaction buffer containing 20 mM Tris-HCl (pH 6.8) and 50 mM NaCl, in a final reaction volume of 120 µL. Reactions were incubated for 5 h at 37°C. Following incubation, insoluble peptidoglycan was removed by centrifugation at 20,000 × *g* for 10 min. The absorbance of soluble RBB-containing peptidoglycan fragments in the resulting supernatant was measured at 595 nm using a Synergy H1 microplate reader (BioTek). Lysozyme (100 µg mL⁻¹) and bovine serum albumin (BSA; 200 µg mL⁻¹) were included as assay controls. For experiments using cell lysates, lysate prepared from *E. coli* carrying the empty-vector (EV) was included as the corresponding lysate control.

#### Growth Experiments and Small-Scale Expression of the T6EC12 proteins

*E. coli* BL21(DE3) cells were transformed with the indicated plasmids and selected on LB agar supplemented with kanamycin. Single colonies were inoculated into LB medium containing kanamycin and 2% glucose and grown overnight at 37 °C with shaking. Overnight cultures were diluted 1:100 into fresh LB medium supplemented with kanamycin and 0.1% glucose and incubated at 37 °C. For growth experiments, cell growth was monitored by measuring the optical density at 600 nm every hour for 12 h. All experiments were performed in biological triplicate. For small-scale expression, cultures were grown to an OD₆₀₀ of 0.5–0.6 and induced with 1 mM IPTG. Following overnight incubation at 30 °C, 0.5-mL culture aliquots were harvested by centrifugation and lysed using BugBuster® Master Mix (Millipore, Cat. No. 71456) supplemented with 1 mM PMSF (Sigma-Aldrich, Israel) and Protease Inhibitor Cocktail Set V (Millipore, Cat. No. 539137). Soluble and insoluble fractions were separated by centrifugation and analyzed by SDS-PAGE and immunoblotting.

#### Construction of Phylogenetic Trees

For T6EC11, T6EC12, and T6EC17, homologous sequences were identified via BLAST (Johnson *et al*, 2008) `using thresholds of ≥40% sequence identity and ≥50% query coverage. Multiple sequence alignments were performed with MUSCLE` (Madeira *et al*, 2024), and phylogenetic trees were reconstructed using FastTree (Price *et al*, 2009) with default parameters. Resulting trees were visualized and annotated using the Interactive Tree of Life (iTOL) (Letunic & Bork, 2024) platform.

## Acknowledgements

We thank Prof. Tommy Kaplan (Hebrew University of Jerusalem) for his advice during the algorithm development. We are grateful to all members of the Levy lab for discussions and comments.

## Data Availability

All the code used in analysis is available in: https://github.com/AvitalAkermanArad/T6SS_ML_Project. Tables EV1 and EV2 are too large to be kept at the supplementary information file. They are stored in Zenodo: https://zenodo.org/records/22370734

## Author Contribution

AAA, NT, and AL designed the study and wrote the manuscript. AAA and AD designed the XGBoost algorithm. YC and YOS are responsible for microscopy analysis with help from GM. AF, SB, MS, and AAA, TF, and RF are responsible for drop assays and biochemical assays.

## Bibliography

Abby SS, Cury J, Guglielmini J, Néron B, Touchon M & Rocha EPC (2016) Identification of protein secretion systems in bacterial genomes. Sci Rep 6: 23080

Abramson J, Adler J, Dunger J, Evans R, Green T, Pritzel A, Ronneberger O, Willmore L, Ballard AJ, Bambrick J, et al (2024) Accurate structure prediction of biomolecular interactions with AlphaFold 3. Nature 630: 493–500

Adeodato PJL & Melo SB (2016) On the equivalence between Kolmogorov-Smirnov and ROC curve metrics for binary classification. arXiv [csAI]

Alcoforado Diniz J, Liu Y-C & Coulthurst SJ (2015) Molecular weaponry: diverse effectors delivered by the Type VI secretion system. Cell Microbiol 17: 1742–1751

Ali J, Yu M, Sung L, Cheung Y & Lai E (2023) A glycine zipper motif is required for the translocation of a T6SS toxic effector into target cells. EMBO Reports 2023 24:6 24: EMBR202356849

Banhos Danneskiold-Samsøe N, Kavi D, Jude KM, Nissen SB, Wat LW, Coassolo L, Zhao M, Santana-Oikawa GA, Broido BB, Garcia KC, et al (2024) AlphaFold2 enables accurate deorphanization of ligands to single-pass receptors. Cell Syst 15: 1046–1060.e3

Benjamini Y & Hochberg Y (1995) Controlling the False Discovery Rate: A Practical and Powerful Approach to Multiple Testing. Journal of the Royal Statistical Society: Series B (Methodological) 57: 289–300

Boyer F, Fichant G, Berthod J, Vandenbrouck Y & Attree I (2009) Dissecting the bacterial type VI secretion system by a genome wide in silico analysis: what can be learned from available microbial genomic resources? BMC Genomics 10: 104

Buchfink B, Reuter K & Drost HG (2021) Sensitive protein alignments at tree-of-life scale using DIAMOND. Nature Methods 2021 18:4 18: 366–368

Campen A, Williams RM, Brown CJ, Meng J, Uversky VN & Dunker AK (2008) TOP-IDP-Scale: A New Amino Acid Scale Measuring Propensity for Intrinsic Disorder. Protein Pept Lett 15: 956–963

Carobbi A, Leo K, Di Nepi S, Bosis E, Salomon D & Sessa G (2024) PIX is an N-terminal delivery domain that defines a class of polymorphic T6SS effectors in Enterobacterales. Cell Rep 43: 114015

Chen T & Guestrin C XGBoost: A Scalable Tree Boosting System.

Chung CT, Niemela SL & Miller RH (1989) One-step preparation of competent Escherichia coli: transformation and storage of bacterial cells in the same solution. Proc Natl Acad Sci U S A 86: 2172–2175

Danov A, Pollin I, Moon E, Ho M, Wilson BA, Papathanos PA, Kaplan T & Levy A (2024a) Identification of novel toxins associated with the extracellular contractile injection system using machine learning. Mol Syst Biol 20: 859–879

Danov A, Segev O, Bograd A, Ben Eliyahu Y, Dotan N, Kaplan T & Levy A (2024b) Toxinome-the bacterial protein toxin database. mBio 15: e0191123

Dong TG, Ho BT, Yoder-Himes DR & Mekalanos JJ (2013) Identification of T6SS-dependent effector and immunity proteins by Tn-seq in Vibrio cholerae. Proceedings of the National Academy of Sciences 110: 2623–2628

Eddy SR (2009) A new generation of homology search tools based on probabilistic inference. Genome Inform 23: 205–211

Eichinger V, Nussbaumer T, Platzer A, Jehl MA, Arnold R & Rattei T (2016) EffectiveDB— updates and novel features for a better annotation of bacterial secreted proteins and Type III, IV, VI secretion systems. Nucleic Acids Res 44: D669–D674

English G, Trunk K, Rao VA, Srikannathasan V, Hunter WN & Coulthurst SJ (2012) New secreted toxins and immunity proteins encoded within the Type VI secretion system gene cluster of Serratia marcescens: Type VI toxins and immunity proteins inSerratia. Mol Microbiol 86: 921–936

Fridman CM, Keppel K, Gerlic M, Bosis E & Salomon D (2020) A comparative genomics methodology reveals a widespread family of membrane-disrupting T6SS effectors. Nat Commun 11: 1085

Fridman CM, Keppel K, Rudenko V, Altuna-Alvarez J, Albesa-Jové D, Bosis E & Salomon D (2025) A new class of type VI secretion system effectors can carry two toxic domains and are recognized through the WHIX motif for export. PLoS Biol 23: e3003053

Fu L, Niu B, Zhu Z, Wu S & Li W (2012) CD-HIT: accelerated for clustering the next-generation sequencing data. Bioinformatics 28: 3150–3152

Galperin MY, Wolf YI, Makarova KS, Alvarez RV, Landsman D & Koonin EV (2021) COG database update: focus on microbial diversity, model organisms, and widespread pathogens. Nucleic Acids Res 49: D274–D281

Gates FL (1933) THE REACTION OF INDIVIDUAL BACTERIA TO IRRADIATION WITH ULTRAVIOLET LIGHT. Science 77: 350

Geller AM, Shalom M, Zlotkin D, Blum N & Levy A (2024) Identification of type VI secretion system effector-immunity pairs using structural bioinformatics. Mol Syst Biol

Geller AM, Zlotkin D & Levy A (2021) Large-scale discovery of candidate type VI secretion effectors with antibacterial activity. bioRxiv: 2021.10.07.463556

González-Magaña A, Altuna J, Queralt-Martín M, Largo E, Velázquez C, Montánchez I, Bernal P, Alcaraz A & Albesa-Jové D (2022) The P. aeruginosa effector Tse5 forms membrane pores disrupting the membrane potential of intoxicated bacteria. Commun Biol 5: 1189

Günther P, Quentin D, Ahmad S, Sachar K, Gatsogiannis C, Whitney JC & Raunser S (2022) Structure of a bacterial Rhs effector exported by the type VI secretion system. PLoS Pathog 18: e1010182

Habich A, Chaves Vargas V, Robinson LA, Allsopp LP & Unterweger D (2025) Distribution of the four type VI secretion systems in Pseudomonas aeruginosa and classification of their core and accessory effectors. Nature Communications 16: 888

Hagan M, Pankov G, Gallegos-Monterrosa R, Williams DJ, Earl C, Buchanan G, Hunter WN & Coulthurst SJ (2023) Rhs NADase effectors and their immunity proteins are exchangeable mediators of inter-bacterial competition in Serratia. Nat Commun 14: 6061

Hersch SJ, Manera K & Dong TG (2020a) Defending against the Type Six Secretion System: beyond Immunity Genes. Cell Rep 33: 108259

Hersch SJ, Watanabe N, Stietz MS, Manera K, Kamal F, Burkinshaw B, Lam L, Pun A, Li M, Savchenko A, et al (2020b) Envelope stress responses defend against type six secretion system attacks independently of immunity proteins. Nat Microbiol 5: 706–714

Hespanhol JT, Sanchez-Limache DE, Nicastro GG, Mead L, Llontop EE, Chagas-Santos G, Farah CS, de Souza RF, Galhardo R da S, Lovering AL, et al (2022) Antibacterial T6SS effectors with a VRR-Nuc domain are structure-specific nucleases. Elife 11: e82437

Hood RD, Singh P, Hsu F, Güvener T, Carl MA, Trinidad RRS, Silverman JM, Ohlson BB, Hicks KG, Plemel RL, et al (2010) A type VI secretion system of Pseudomonas aeruginosa targets a toxin to bacteria. Cell Host Microbe 7: 25–37

Huang A, Lu M, Ling E, Li P & Wang C (2020) A M35 family metalloprotease is required for fungal virulence against insects by inactivating host prophenoloxidases and beyond. Virulence 11: 222–237

Hu Y, Yan M, Zhu Y, Chao H, Li S, Ni Q, Hu Y, Liu E, Liu L, Chen Y, et al (2025) Improved Prediction of Bacterial Type VI Secretion Effector Proteins Using an Integrated Convolutional Neural Network Model Combining N-terminal Signal Sequences, Evolutionary Information and Pre-Trained Protein Language Features. bioRxiv: 2025.03.07.642067–2025.03.07.642067

Jana B, Fridman CM, Bosis E & Salomon D (2019) A modular effector with a DNase domain and a marker for T6SS substrates. Nature Communications 10: 3595

Jiang X, Li H, Ma J, Li H, Ma X, Tang Y, Li J, Chi X, Deng Y, Zeng S, et al (2024) Role of Type VI secretion system in pathogenic remodeling of host gut microbiota during Aeromonas veronii infection. ISME J 18

Jiaweiwang J, Yang B, Leier A, Marquez-Lago TT, Hayashida M, Rocker A, Zhang Y, Akutsu T, Chou KC, Strugnell RA, et al (2018) Bastion6: a bioinformatics approach for accurate prediction of type VI secreted effectors. Bioinformatics 34: 2546–2555

Johnson M, Zaretskaya I, Raytselis Y, Merezhuk Y, McGinnis S & Madden TL (2008) NCBI BLAST: a better web interface. Nucleic Acids Res 36: W5–9

van Kempen M, Kim SS, Tumescheit C, Mirdita M, Lee J, Gilchrist CLM, Söding J & Steinegger M (2023) Fast and accurate protein structure search with Foldseek. bioRxiv: 2022.02.07.479398–2022.02.07.479398

Koskiniemi S, Lamoureux JG, Nikolakakis KC, De Roodenbeke CTK, Kaplan MD, Low DA & Hayes CS (2013) Rhs proteins from diverse bacteria mediate intercellular competition. Proceedings of the National Academy of Sciences 110: 7032–7037

Kudryashev M, Wang RY-R, Brackmann M, Scherer S, Maier T, Baker D, DiMaio F, Stahlberg H, Egelman EH & Basler M (2015) Structure of the type VI secretion system contractile sheath. Cell 160: 952–962

Kyte J & Doolittle RF (1982) A simple method for displaying the hydropathic character of a protein. J Mol Biol 157: 105–132

LaCourse KD, Peterson SB, Kulasekara HD, Radey MC, Kim J & Mougous JD (2018) Conditional toxicity and synergy drive diversity among antibacterial effectors. Nat Microbiol 3: 440–446

Lederberg J (1956) BACTERIAL PROTOPLASTS INDUCED BY PENICILLIN. Proc Natl Acad Sci U S A 42: 574–577

Le N-H, Pinedo V, Lopez J, Cava F & Feldman MF (2021) Killing of Gram-negative and Gram-positive bacteria by a bifunctional cell wall-targeting T6SS effector. Proc Natl Acad Sci U S A 118: e2106555118

Letunic I & Bork P (2024) Interactive Tree of Life (iTOL) v6: recent updates to the phylogenetic tree display and annotation tool. Nucleic Acids Res 52: W78–W82

Levy A, Salas Gonzalez I, Mittelviefhaus M, Clingenpeel S, Herrera Paredes S, Miao J, Wang K, Devescovi G, Stillman K, Monteiro F, et al (2017) Genomic features of bacterial adaptation to plants. Nature Genetics 2017 50:1 50: 138–150

Lewis JM, Deveson Lucas D, Harper M & Boyce JD (2019) Systematic identification and analysis of acinetobacter baumannii type vi secretion system effector and immunity components. Front Microbiol 10: 489615–489615

Liang X, Moore R, Wilton M, Wong MJQ, Lam L & Dong TG (2015) Identification of divergent type VI secretion effectors using a conserved chaperone domain. Proc Natl Acad Sci U S A 112: 9106–9111

Li J, Yao Y, Xu HH, Hao L, Deng Z, Rajakumar K & Ou HY (2015) SecReT6: A web-based resource for type VI secretion systems found in bacteria. Environ Microbiol 17: 2196–2202

Lin J-S, Ma L-S & Lai E-M (2013) Systematic dissection of the agrobacterium type VI secretion system reveals machinery and secreted components for subcomplex formation. PLoS One 8: e67647

Lin J, Zhang W, Cheng J, Yang X, Zhu K, Wang Y, Wei G, Qian PY, Luo ZQ & Shen X (2017) A Pseudomonas T6SS effector recruits PQS-containing outer membrane vesicles for iron acquisition. Nature Communications 2017 8:1 8: 14888

MacIntyre DL, Miyata ST, Kitaoka M & Pukatzki S (2010) The Vibrio cholerae type VI secretion system displays antimicrobial properties. Proc Natl Acad Sci U S A 107: 19520–19524

Madeira F, Madhusoodanan N, Lee J, Eusebi A, Niewielska A, Tivey ARN, Lopez R & Butcher S (2024) The EMBL-EBI Job Dispatcher sequence analysis tools framework in 2024. Nucleic Acids Res 52: W521–W525

Mamou G, Corona F, Cohen-Khait R, Housden NG, Yeung V, Sun D, Sridhar P, Pazos M, Knowles TJ, Kleanthous C, et al (2022) Peptidoglycan maturation controls outer membrane protein assembly. Nature 606: 953–959

Markowitz VM, Chen IMA, Palaniappan K, Chu K, Szeto E, Grechkin Y, Ratner A, Jacob B, Huang J, Williams P, et al (2012) IMG: the integrated microbial genomes database and comparative analysis system. Nucleic Acids Res 40: D115–D122

Mistry J, Chuguransky S, Williams L, Qureshi M, Salazar GA, Sonnhammer ELL, Tosatto SCE, Paladin L, Raj S, Richardson LJ, et al (2021) Pfam: The protein families database in 2021. Nucleic Acids Res 49: D412–D419

Mok BY, de Moraes MH, Zeng J, Bosch DE, Kotrys AV, Raguram A, Hsu F, Radey MC, Peterson SB, Mootha VK, et al (2020) A bacterial cytidine deaminase toxin enables CRISPR-free mitochondrial base editing. Nature 583: 631–637

Mondino S, Schmidt S & Buchrieser C (2020) Molecular mimicry: A paradigm of host-microbe coevolution illustrated by legionella. MBio 11: 1–19

More AS & Rana DP (2017) Review of random forest classification techniques to resolve data imbalance. In 2017 1st International Conference on Intelligent Systems and Information Management (ICISIM) pp 72–78. IEEE

Nachmias N, Dotan N, Rocha MC, Fraenkel R, Detert K, Kluzek M, Shalom M, Cheskis S, Peedikayil-Kurien S, Meitav G, et al (2024) Systematic discovery of antibacterial and antifungal bacterial toxins. Nature Microbiology 2024 9:11 9: 3041–3058

Nicastro GG, Sibinelli-Sousa S, Hespanhol JT, Santos TWC, Munoz JP, Santos RS, Perez-Sepulveda BM, Miyamoto S, Aravind L, de Souza RF, et al (2026) Systematic identification of Salmonella T6SS effectors uncovers diverse new families and lipid-targeting activities. PLoS Biol 24: e3003680

Nolan LM, Cain AK, Clamens T, Furniss RCD, Manoli E, Sainz-Polo MA, Dougan G, Albesa-Jové D, Parkhill J, Mavridou DAI, et al (2021) Identification of Tse8 as a Type VI secretion system toxin from Pseudomonas aeruginosa that targets the bacterial transamidosome to inhibit protein synthesis in prey cells. Nat Microbiol 6: 1199–1210

Paracuellos P, Bexter A, Patkowski JB, Kelly SD, Omelchenko O, Macé K, Ilangovan A, Subramoni S, Whitney JC, Filloux A, et al (2026) Molecular basis of type VI secretion system effector loading. Nat Microbiol 11: 1982–1994

Paysan-Lafosse T, Blum M, Chuguransky S, Grego T, Pinto BL, Salazar GA, Bileschi ML, Bork P, Bridge A, Colwell L, et al (2023) InterPro in 2022. Nucleic Acids Res 51: D418–D427

Price MN, Dehal PS & Arkin AP (2009) FastTree: computing large minimum evolution trees with profiles instead of a distance matrix. Mol Biol Evol 26: 1641–1650

Pukatzki S, Ma AT, Revel AT, Sturtevant D & Mekalanos JJ (2007) Type VI secretion system translocates a phage tail spike-like protein into target cells where it cross-links actin. Proc Natl Acad Sci U S A 104: 15508–15513

Pukatzki S, Ma AT, Sturtevant D, Krastins B, Sarracino D, Nelson WC, Heidelberg JF & Mekalanos JJ (2006) Identification of a conserved bacterial protein secretion system in Vibrio cholerae using the Dictyostelium host model system. Proc Natl Acad Sci U S A 103: 1528– 1533

Ray A, Schwartz N, de Souza Santos M, Zhang J, Orth K & Salomon D (2017) Type VI secretion system MIX-effectors carry both antibacterial and anti-eukaryotic activities. EMBO Rep 18: 1978–1990

Rehman S, Sa-Pessoa J, Buckley C, Lee RL, Wojtania K, Lee A, Lancaster R, Augustine J, Barabas P, Ross C, et al (2026) Klebsiella pneumoniae inhibits vasodilation through capsule and T6SS-dependent pathways. Nat Microbiol

Rocha VD, Dal’Sasso TCS, Costa MDBL & de Oliveira LO (2026) Evolutionary history and diversification of M35 metalloproteases in Dothideomycetes: A phylogenomic overview and case study in Corynespora cassiicola. Curr Microbiol 83

Salomon D, Kinch LN, Trudgian DC, Guo X, Klimko JA, Grishin NV, Mirzaei H & Orth K (2014) Marker for type VI secretion system effectors. Proc Natl Acad Sci U S A 111: 9271–9276

Salomon D, Klimko JA, Trudgian DC, Kinch LN, Grishin NV, Mirzaei H & Orth K (2015) Type VI secretion system toxins horizontally shared between marine bacteria. PLoS Pathog 11: e1005128

Sana TG, Baumann C, Merdes A, Soscia C, Rattei T, Hachani A, Jones C, Bennett KL, Filloux A, Superti-Furga G, et al (2015) Internalization of Pseudomonas aeruginosa strain PAO1 into epithelial cells is promoted by interaction of a T6SS effector with the microtubule network. MBio 6: e00712

Sánchez-Gorostiaga A, Palacios P, Martínez-Arteaga R, Sánchez M, Casanova M & Vicente M (2016) Life without Division: Physiology of Escherichia coli FtsZ-Deprived Filaments. mBio 7

Sen R, Nayak L & De RK (2019) PyPredT6: A python-based prediction tool for identification of Type VI effector proteins. https://doi.org/101142/S0219720019500197 17

Shidore T, Dinse T, Öhrlein J, Becker A & Reinhold-Hurek B (2012) Transcriptomic analysis of responses to exudates reveal genes required for rhizosphere competence of the endophyte Azoarcus sp. strain BH72: Endophyte response to exudates. Environ Microbiol 14: 2775– 2787

Si M, Zhao C, Burkinshaw B, Zhang B, Wei D, Wang Y, Dong TG & Shen X (2017) Manganese scavenging and oxidative stress response mediated by type VI secretion system in Burkholderia thailandensis. Proceedings of the National Academy of Sciences 114: E2233– E2242

Steinegger M & Söding J (2017) MMseqs2 enables sensitive protein sequence searching for the analysis of massive data sets. Nat Biotechnol 35: 1026–1028

Tallarida RJ & Murray RB (1987) Chi-Square Test. Manual of Pharmacologic Calculations: 140– 142

Ting S-Y, Bosch DE, Mangiameli SM, Radey MC, Huang S, Park Y-J, Kelly KA, Filip SK, Goo YA, Eng JK, et al (2018) Bifunctional Immunity Proteins Protect Bacteria against FtsZ-Targeting ADP-Ribosylating Toxins. Cell 175: 1380–1392.e14

Unni R, Pintor KL, Diepold A & Unterweger D (2022) Presence and absence of type VI secretion systems in bacteria: This article is part of the Bacterial Cell Envelopes collection. Microbiology 168

Verster AJ, Ross BD, Radey MC, Bao Y, Goodman AL, Mougous JD & Borenstein E (2017) The Landscape of Type VI Secretion across Human Gut Microbiomes Reveals Its Role in Community Composition. Cell Host Microbe 22: 411–419.e4

Wang C, Chen M, Shao Y, Jiang M, Li Q, Chen L, Wu Y, Cen S, Waterfield NR, Yang J, et al (2023) Genome wide analysis revealed conserved domains involved in the effector discrimination of bacterial type VI secretion system. Communications Biology 6: 1195

Wang J, Li J, Hou Y, Dai W, Xie R, Marquez-Lago TT, Leier A, Zhou T, Torres V, Hay I, et al (2021) BastionHub: a universal platform for integrating and analyzing substrates secreted by Gram-negative bacteria. Nucleic Acids Res 49: D651–D659

Whitney JC, Chou S, Russell AB, Biboy J, Gardiner TE, Ferrin MA, Brittnacher M, Vollmer W & Mougous JD (2013) Identification, structure, and function of a novel type VI secretion peptidoglycan glycoside hydrolase effector-immunity pair. J Biol Chem 288: 26616–26624

Yang M, Derbyshire MK, Yamashita RA & Marchler-Bauer A (2020) NCBI’s Conserved Domain Database and tools for protein domain analysis. Curr Protoc Bioinformatics 69: e90

Zhu L, Xu L, Wang C, Li C, Li M, Liu Q, Wang X, Yang W, Pan D, Hu L, et al (2021) T6SS translocates a micropeptide to suppress STING-mediated innate immunity by sequestering manganese. Proc Natl Acad Sci U S A 118: e2103526118–e2103526118

